# Telmisartan impedes JEV infection predominately via AT1/PPARγ axis

**DOI:** 10.1101/2024.10.29.620930

**Authors:** Ankita Datey, Sanchari Chatterjee, Soumyajit Ghosh, P Sanjai Kumar, Saikat De, Udvas Ghorai, Bharat Bhusan Subudhi, Soma Chattopadhyay

## Abstract

Japanese encephalitis, caused by Japanese encephalitis virus (JEV) is a vector born disease for which no specific therapeutics are available yet. Binding of angiotensin II (Ang II) to angiotensin II type 1 (AT1) receptor induces release of inflammatory cytokines associated with viral encephalitis. Accordingly, Ang II receptor blockers (ARBs) have been proposed to manage encephalitis. Since Telmisartan (TM, antagonist of AT1 and agonist of PPARγ) has relatively better brain access than other ARBs, this investigation aims to evaluate its anti-JEV efficacy *in vitro* and *in vivo.* TM reduced JEV titer, RNA and protein (NS3) significantly in the BHK-21 cells with IC_50_ of 24.68μM and CC_50_ of >350 μM (Selectivity Index >14.18) indicating its potential of repurposing against JEV. The anti-JEV efficacy of TM was further observed in macrophages (RAW264.7) and neuronal (SH-SY5Y) cells. Interestingly, the viral load was reduced significantly in pre, co and post-treatment conditions of TM, however most efficiently (80%) in post treatment. In presence of GW (PPARγ antagonist) and AG (AT1 agonist), viral infection was increased remarkably while AT1 was upregulated and PPARγ was downregulated. Whereas, TM treatment reversed the levels of AT1 and PPARγ during infection. Further, reduction of inflammatory markers like p-IRF-3, COX-2 and p-NF-κB was observed after TM treatment in RAW264.7 cells suggesting its immunomodulation through the AT1/PPARγ axis. Finally, the anti-JEV potential of TM was validated in mice model through the reduction of disease score, viral protein and histological changes. Thus, the preclinical efficacy of TM suggests its suitability for repurposing against JEV.

## 1. Introduction

JEV (Japanese encephalitis virus), a mosquito born virus belongs to family *Flaviviridae.* It comprises of single stranded positive sense RNA and size of the genome is around 11kb. Its genome is divided into three structural proteins, namely capsid (C), pre-membrane (prM), and envelope (E), and seven non-structural proteins (NS1, NS2A, NS2B, NS3, NS4A, NS4B, and NS5) (Guo et al., 2020). It primarily causes encephalitis, affecting children upto 14 years of age whereas, adults develop immunity after infection (Campbell et al., 2011). JEV is transmitted by *Culex* species of mosquitoes mainly *Culex vishnuii* and *Culex tritaeniorhynchus*. Pigs act as an amplifier host, whereas bats along with Ardeid birds (herons and egrets) act as a primary reservoir host for JEV. With prolonged viraemia, mosquitoe vectors transmit JEV to humans who are considered as dead-end host. The common sign and symptoms include fever, nausea, headache, movement disorders, whereas severe condition results in permanent neurological manifestation which often leads to mortality (Simon et al., 2024). Although incidences of encephalitis are limited to approximately 1% of JEV-infected population, higher rates (20-30%) of mortality is associated with individuals developing JE. Further, 50% of patient who survive JE, develop neuropsychiatric sequelae (Joe et al., 2022). Globally, 30000-50000 incidences of JEV are reported each year mostly from twenty-four countries in South-East Asia and the Western Pacific where it is endemic (Simon et al., 2024). Far away countries including Australia have also been identified as risk areas by Centre for Disease Control (CDC), USA.

Vaccines are available for JEV. However, high cost, multiple doses, emerging genotypes and poorly coordinated JEV control programs are some of the challenges to its effectiveness in preventing JE (Vannice et al., 2021). Further, enzootic cycle of JEV makes it difficult to completely eradicate. Thus, there is a need for its therapeutic management. Although this has drawn attention for antiviral research, no specific therapy is available for application. Drug repurposing is a better alternative to new drug development to minimize cost and time for developing antiviral strategy. Accordingly, several drugs such as minocycline, interferon, ribavirin, immunoglobulin, dexamethasone and acyclovir have been investigated against JEV (Ajibowo et al., 2021). Out of these, only minocycline showed statistically significant decrease. However, till date no specific antiviral drug is available against JEV that necessitates exploring alternative approaches (Joe et al., 2022).

Considering that encephalitis is the major reason of mortality and morbidity associated with JEV infection, it is worthwhile to explore drugs capable of reducing neuroinflammation for anti-JEV therapy. Because of the involvement of local renin–angiotensin system (RAS) in the neuroinflammation, inhibiting RAS has been documented as a strategy to minimize encephalitis (Subudhi & Sahu, 2023). Binding of angiotensin II (Ang II) to angiotensin II type1 (AT1) receptor is known to induce release of inflammatory cytokines associated with viral encephalitis (Chabrashvili et al., 2003). Accordingly, angiotensin II receptor blockers (ARBs) have been proposed to manage encephalitis. However, majority of the approved ARBs have poor access to brain because of inability to cross blood brain barrier (BBB). Nevertheless, increasing their brain access has been shown to enhance protection against neuroinflammation (Subudhi et al., 2018). Unlike most of these ARBs, telmisartan (TM) has relatively better brain access and selectively blocks AT1R activation (proinflammatory axis) without hindering signaling cascade of AT2R (anti-inflammatory axis) (Blakely et al., 2016). Further, it is also a partial agonist of peroxisome proliferator activated receptor gama (PPAR)γ. Down-regulation of PPARγ is an important marker of viral infection-induced inflammation in brain (Layrolle et al., 2021). Hence, TM can be useful to manage encephalitis associated with JEV. Interestingly, our group has earlier shown antiviral properties of TM against Chikungunya virus (CHIKV) partly mediated through modulation of PPARγ and AT1 (De et al., 2022). Few studies have also shown antiviral properties of AT1 blockers against Venezuelan equine encephalitis virus (VEEV) and flavivirus infections like Dengue (Bermúdez et al., 2015; Hernández-Fonseca et al., 2015) . Since there are no such reports available against JEV till now, the current study aims to evaluate the anti-viral efficacy of TM against JEV *in vitro* and *in vivo*.

It was observed that TM was able to reduce the viral titer by 90% *in vitro.* To better understand the mechanism of action, GW (PPARγ antagonist) and AG (AT1 agonist) were used. In presence of GW and AG, viral infection was enhanced suggesting role of AT1/PPARγ in JEV infection. Further, detailed mechanism of action of TM revealed downregulation of p-IRF-3 (phospho - interferon regulatory factor 3), COX-2 (cyclooxygenase-2) and p-NF-κB (phospho -nuclear factor kappa-light-chain-enhancer of activated B cells) responsible for inflammation. It was further validated by *in vivo* studies in Balb/c mice model where the JEV-NS3 protein was reduced to 60% approximately in TM treated mice brain as compared to infection. Thus, the above findings reveal that TM might be repurposed efficiently to restrict JEV infection.

## 2. Results

### 2.1. Telmisartan hinders JEV infection *in vitro*

In order to understand the cytotoxicity of TM in BHK21 cells, MTT (3-(4,5-dimethylthiazol-2-yl)-2,5-diphenyltetrazolium bromide) assay was performed with different concentrations of this molecule and 50% cytotoxicity (CC_50_) was determined. It was observed that the drug was non-cytotoxic till 350µM concentration (Fig. 1A). To determine the antiviral efficacy of TM, BHK-21cells (Baby Hamster Kidney-21) were infected with JEV at an MOI of 0.1 and TM was administered post infection at varying concentrations (25, 50 and 75µM). The cell culture supernatant and cells were harvested at 24 hours post infection (hpi). It was observed that there was a significant dose dependent reduction of 25%, 50% and 75% in viral copy no/mL and 45, 52 and 70% reduction in JEV-NS3 (JEV-Nonstructural protein3) protein level approximately (Fig. 1B, C and D). The data was further supported by the confocal analysis (Fig. 1E and F). Further, the viral titer showed remarkable reduction of 80, 90 and 95% with increasing concentrations (25, 50 and 75µM)) of TM respectively (Fig. 1G). The IC_50_ (Inhibitory concentration 50) of TM was found to be 24.68μM (Fig. 2A) and the selectivity index was observed to be 14.18 (Supplementary Table.S2). Further, to investigate the antiviral activity of JEV in physiologically relevant cell lines such as RAW264.7 and SH-SY5Y cells, JEV infection was carried out at MOI of 5 and 0.1 respectively. It was observed that JEV-E (Envelope) gene expression was decreased upto 60%, 70% and 80% at 25, 50 and 75µM concentrations of TM respectively in RAW264.7 cell through qRT-PCR. Whereas up to 10%, 25% and 75% reduction was observed at 25, 50 and 75µM concentrations of TM in SH-SY5Y cells respectively (Supplementary Fig. 1A and D). Similarly, INF-β (Interferon-β) gene expression and iNOS (Inducible nitric oxide synthase) levels in RAW264.7 cells were also down regulated remarkably with TM treatment (Supplementary Fig. 1B and C). Moreover, expression of Cas3 (Caspase3) gene in SH-SY5Y cells was also downregulated significantly in TM treated samples (Supplementary Fig. 1E). Thus, the data suggest that TM efficiently inhibits the JEV infection *in vitro* in various cell lines and also downregulates levels of inflammatory cytokines and apoptotic markers.

**Fig. 1.**
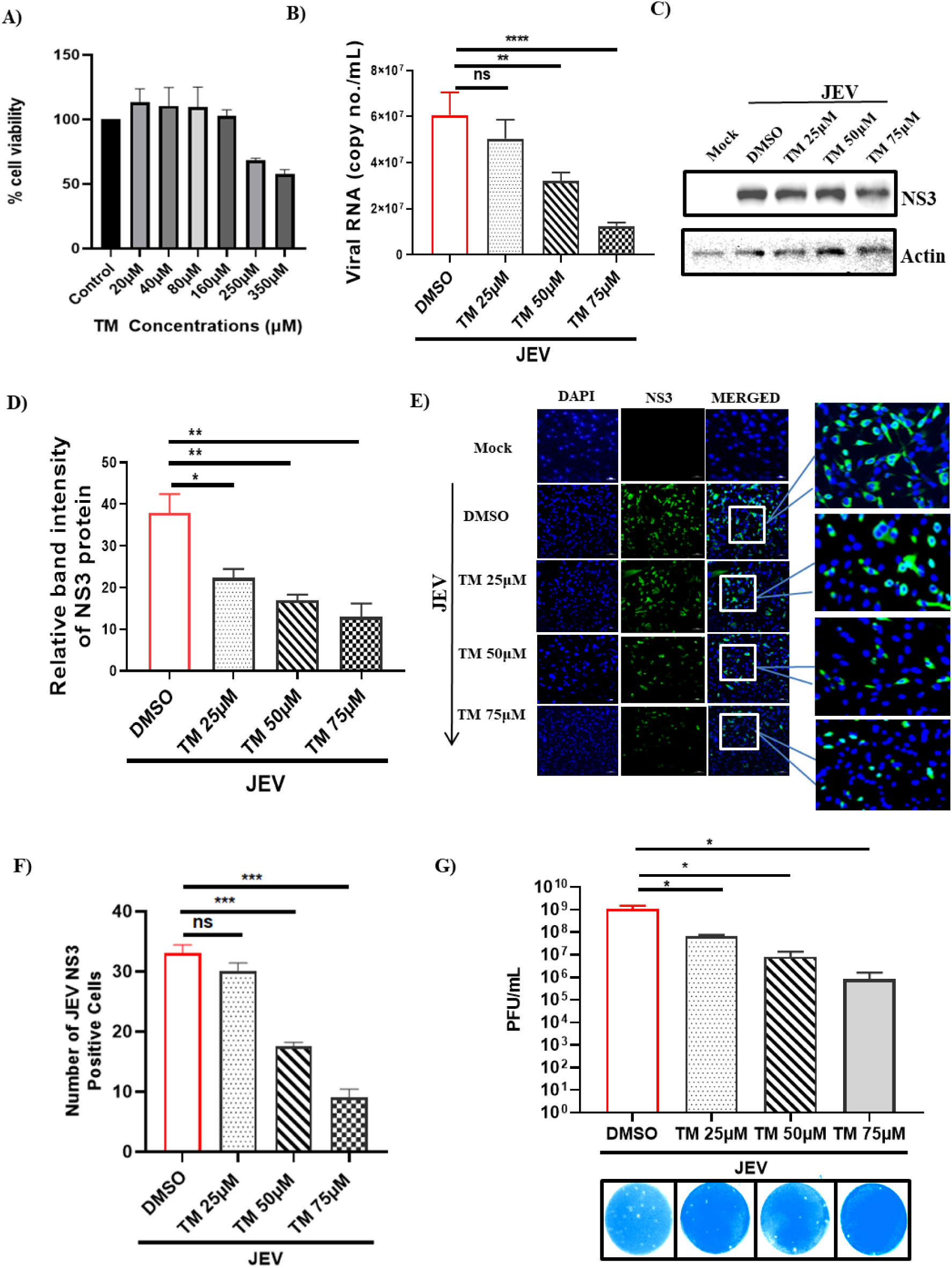
TM possesses anti-JEV activity: BHK-21 cells were infected with JEV at MOI 0.1 and TM was administered at varying concentrations (25, 50 and 75µM) post infection. The supernatant and pellets were harvested after 24hpi and subjected to qRT-PCR and Western blot respectively. (A) Bar diagram showing viability of cells in presence of TM. (B) Bar diagram representing viral RNA copy no./mL with increasing TM concentration. (C)Western blot image depicting the level of the JEV-NS3 protein in samples treated with varying concentration of TM. (D) Bar diagram showing relative band intensities of the JEV-NS3 protein. (E) Confocal image showing mock, infected and TM treated cells, stained with NS3 antibody and nuclei were counterstained with DAPI (Scale bar 100µM). (F) Bar diagram representing percentage of JEV-NS3 positive cells. (G) Bar diagram showing the viral titers (PFU/mL) of different samples. Lower panel shows the images of plaque assay plates. The analysis was carried out using the one-way ANOVA test. *, *P* ≤ 0.05; **, *P* ≤ 0.01; ***, *P* ≤ 0.001; and ****, *P* ≤ 0.0001 were considered statistically significant and ns, not significant (n=3).

**Fig. 2.**
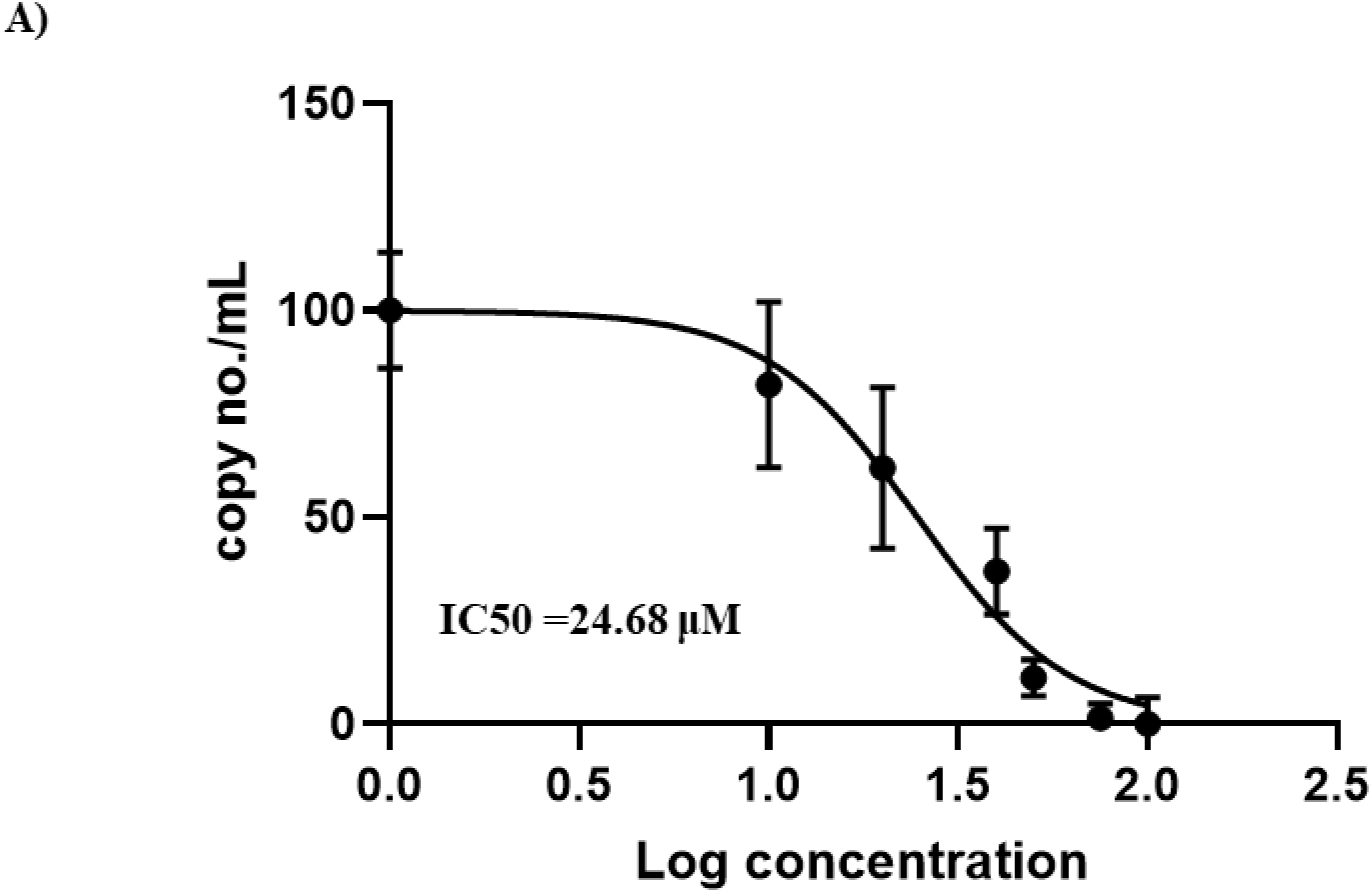
Determining the IC_50_ value of TM: For calculating IC_50_ value of TM, BHK-21 cells were infected with JEV (MOI 0.1) and different concentrations of TM (10, 20, 30, 40, 50, 75, 100, 125 and 150µM) were added post infection. The supernatant was collected and qRT-PCR was performed to determine the viral RNA as copy no./mL. (A) Graph depicting IC_50_ of TM, where the X-axis represents the logarithmic value of varying TM concentrations and the Y-axis represents the percentage of viral RNA in terms of copy no./mL.

### 2.2. Pre, co and post-treatment of TM can abrogate JEV infection

To identify at which stage of the viral life cycle TM (50µM) can inhibit JEV infection, the drug was administered in pre, co and post infection conditions as mentioned above. It was observed that TM efficiently reduced the viral titer in all the three conditions to 50%, 60% and 80% respectively but most efficiently in post condition (Fig. 3A). Further, to find out whether TM has direct impact on JEV viral particles, virus inactivation or virucidal assay was performed. There was significant inhibition (30% approx) in viral titer in drug treated sample as that of infection control (Fig. 3B). Thus, TM might directly impair JEV particles which may interfere in attachment and/entry. For better understanding the effect of TM in entry or attachment, infection was carried out with JEV in presence or absence of TM. After the infection was over, unbound virus particles as well as the new particles formed after 24 hpi were collected and the viral copy no./mL and viral titers (PFU/mL) were determined. Interestingly, it was observed that viral copy no. as well as titer of unbound particles in presence of TM was less as compared to unbound virus particles in absence of TM. Further, the new viral particles formed after 24hpi was remarkably reduced in presence of TM (viral copy no and titer) as that of infection control (Fig. 3C and D). Together, the data suggest that TM interferes with JEV infection in different stages of its life cycle.

**Fig. 3.**
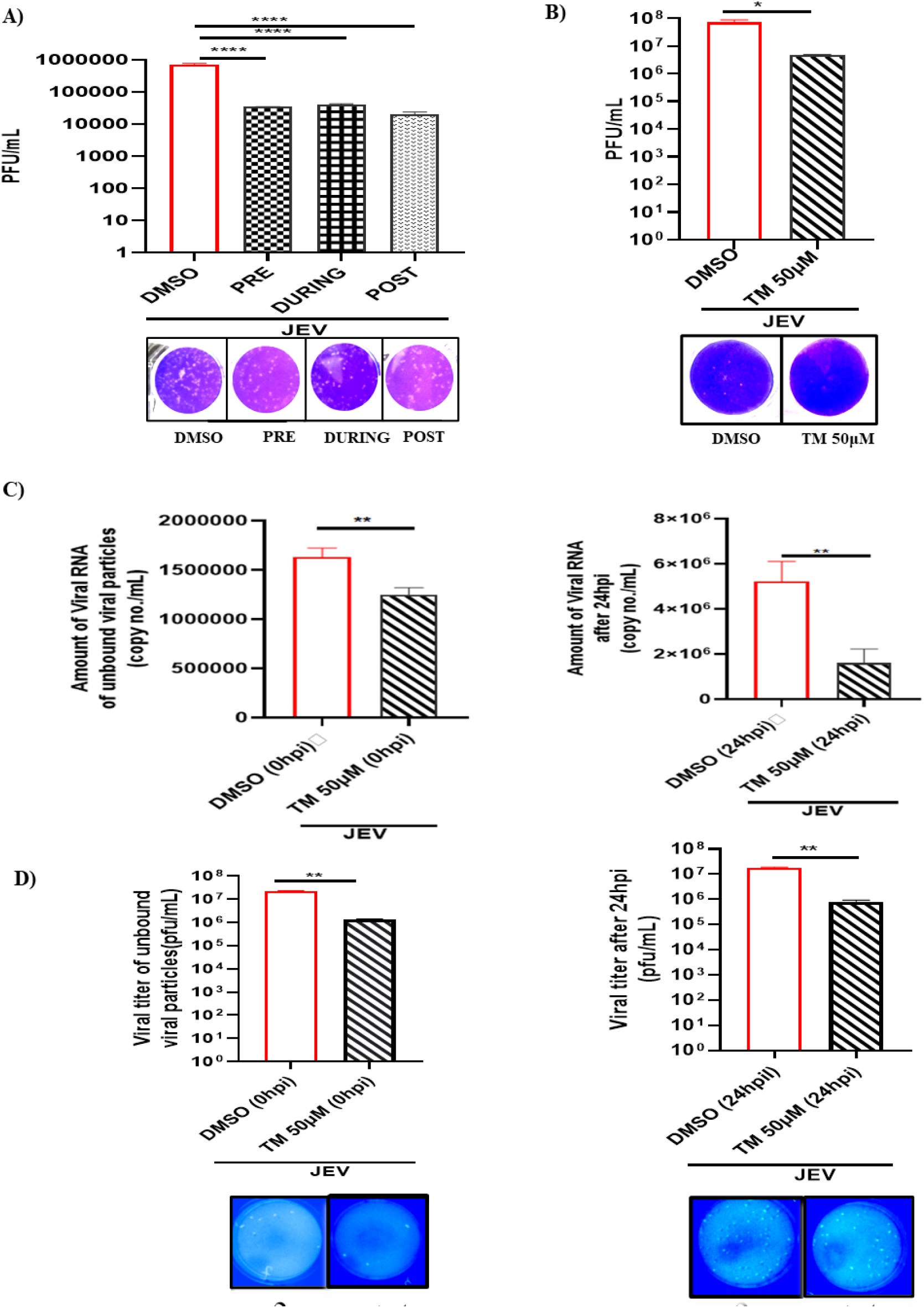
TM inhibits JEV infection in pre, co and post treatment: BHK-21 cells were treated with TM (50µM) for 3hrs prior to infection (MOI 0.1) in pre-condition; in during condition TM was given along with virus whereas in post condition TM was administered post infection. The supernatant was collected after 24 hpi and plaque assay was performed. (A) Bar diagram representing viral titers in all the three conditions. Lower panel represents the images of plaque assay plates. (B) Bar diagram representing the virucidal activity of TM in absence or presence of drug. Lower region shows the image of plaque assay plates. (C) Bar diagram representing the viral RNA copy no./mL in different samples (unbound virus particle; left panel whereas viral particles formed after 24hpi; right panel). (D) Bar diagram showing the viral titers in various samples (unbound virus particle forming plaques; left panel whereas viral particles collected at 24hpi generating plaques; right panel). Lower panel depicts the images of plaque assay plates. Data shown is the repetition of three individual experiments with standard mean represented as ± SD. The one-way ANOVA and unpaired two-tailed Student’s *t* test were performed for all the experiments. *, *P* ≤ 0.05; **, *P* ≤ 0.01; ***, *P* ≤ 0.001 were considered statistically significant.

### 2.3. TM reduces JEV titer through AT1/PPARγ pathway

As TM is the antagonist of AT1 and agonist of PPARγ, antagonist of PPARγ (GW) and agonist of AT1 receptor (AG) were utilized to understand the mechanism of action of TM in JEV infection. First their cytotoxicity was determined. GW and AG were found to be non-toxic till 80µM and 100µM concentrations respectively (Fig. 4A and 5A). The BHK-21 cells were pre treated for 3 hrs with increasing concentrations of GW (1, 5, 10 and 20µM) or AG (10, 20, 30 and 40µM). Post-infection, again the compounds were added and samples (supernatant and cells) were harvested after 24 hpi. It was observed that the viral copy number, JEV-NS3 protein level and viral titer were significantly increased with increasing concentrations of GW and AG (Fig. 4B, C, D, E and 5B, C, D, E). Similar observations were also noticed by the confocal microscopy (Supplementary Fig. 2S A, B, C and D). Further, the FACS data revealed significant increase in expression of the JEV-NS3 protein with increased concentrations of GW (Fig. 4F and G).

**Fig. 4.**
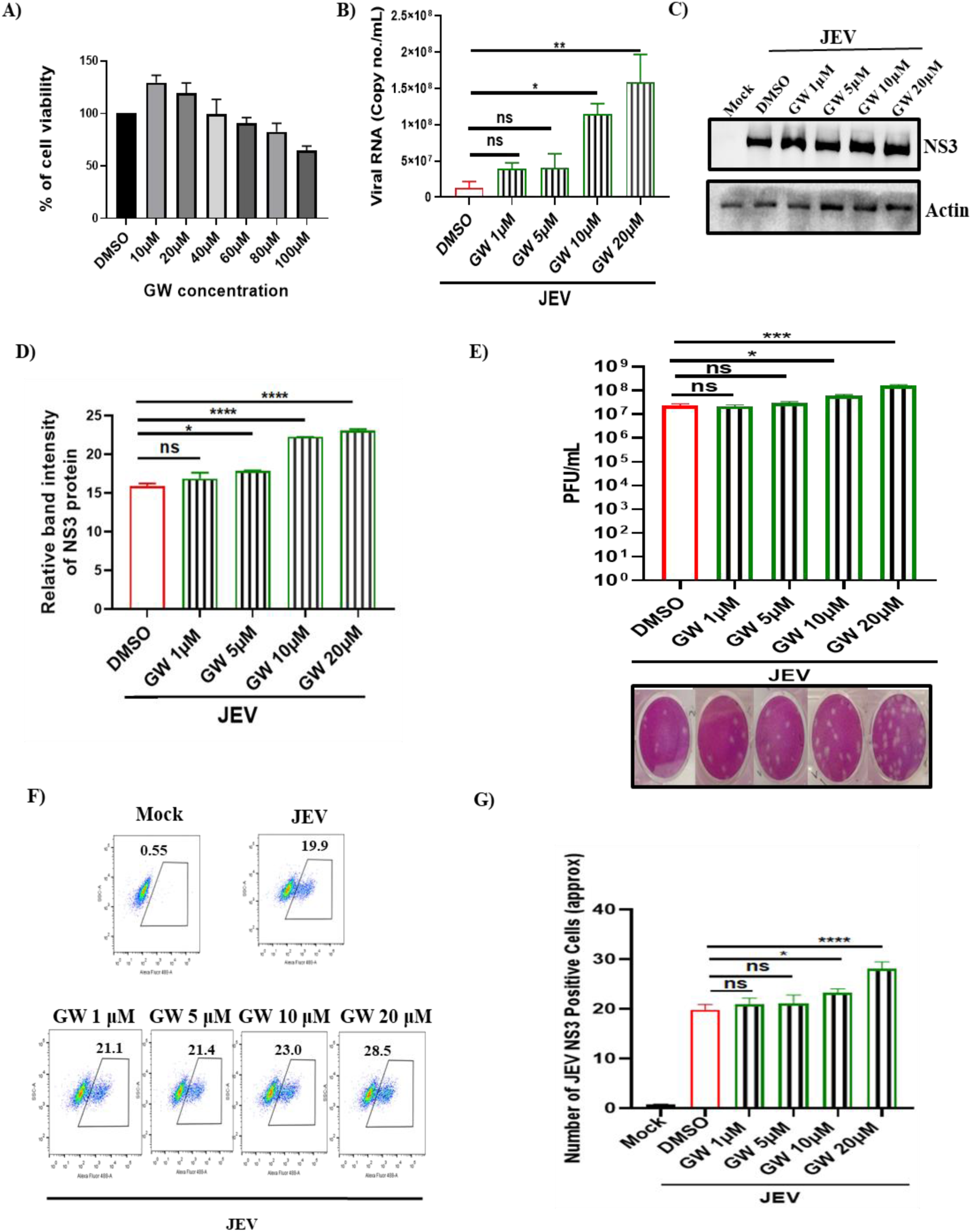
GW (PPARγ antagonist) enhances viral load: JEV infection was carried out in GW (1, 5, 10 and 20µM) pretreated (3hrs) BHK-21 cells and GW was administered again after infection. The supernatant and pellets were collected at 24 hpi. (A) Bar graph representing the cytotoxicity of GW. (B) Bar diagram representing the viral RNA in terms of copy no./mL in different samples. (C) Western blot image representing the JEV-NS3 protein level in different samples. Actin was used as a loading control. (D) Bar diagram depicting the relative band intensity of JEV-NS3 protein. (E) Bar diagram represents the viral plaques in various samples. Lower panel represents the plaque assay plate image. (F and G) Dot plot analysis and bar diagram showing the number of JEV-NS3 positive BHK-21 cells. All the experiments were carried out in triplicates and calculation was done using the one-way ANOVA test. *, *P* ≤ 0.05; **, *P* ≤ 0.01; and ****, *P* ≤ 0.0001 were considered statistically significant. ns, not significant

**Fig. 5.**
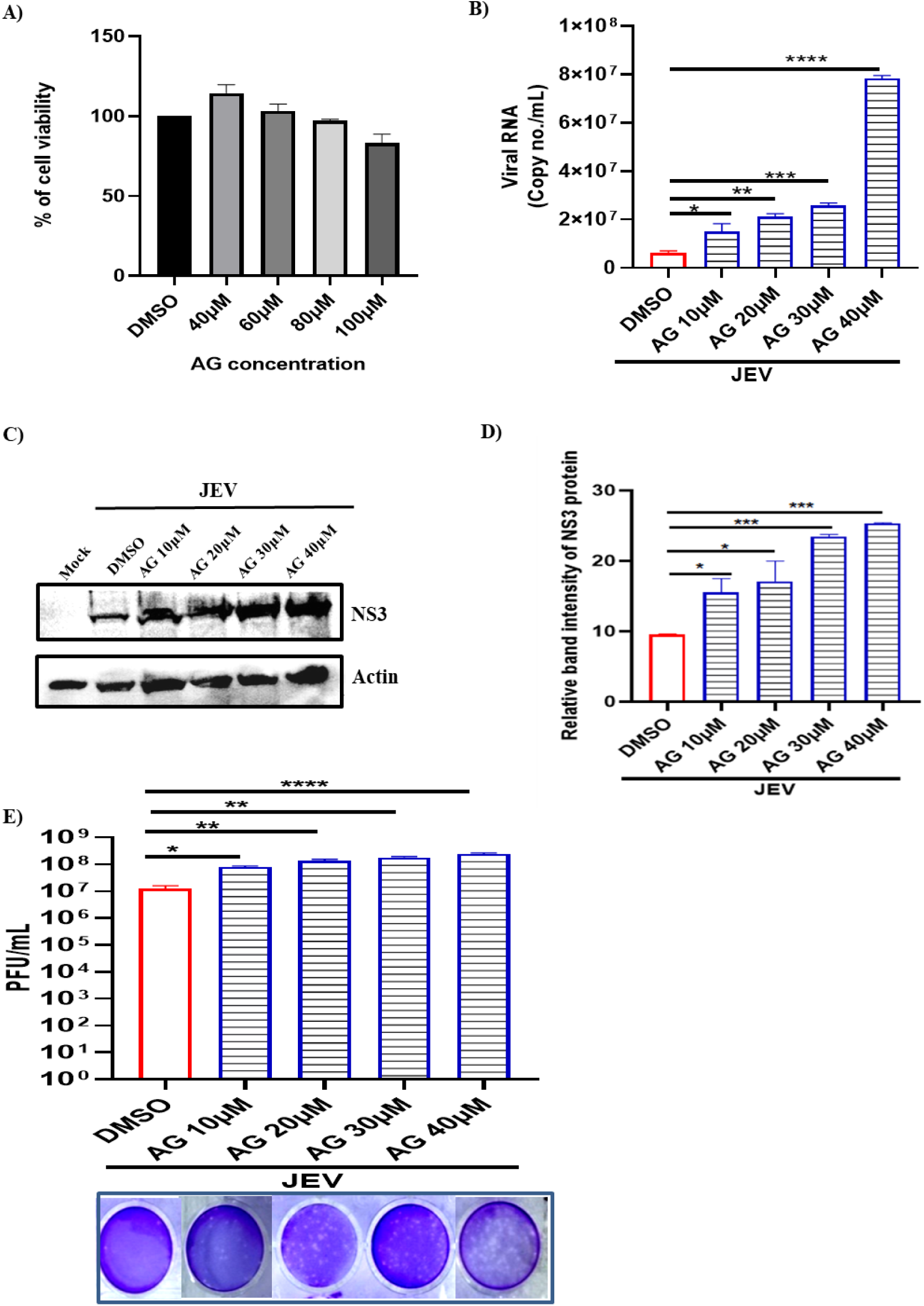
AG, a AT1 agonist augments viral infection: The BHK-21 cells were pretreated for 3hrs with varying AG concentrations (10, 20, 30 and 40µM). Later, JEV infection (MOI 0.1) was given to the cells and AG was administered again post infection. After 24hpi, supernatant and pellet were harvested. (A) Bar graph showing percentage of viable cells. (B) Bar graph indicating the viral RNA in terms of copy no./mL in various samples. (C) Western Blot image showing the band intensity of the JEV-NS3 protein in various samples. Actin was used as a loading control. (D) Bar diagram depicting the relative band intensity of the JEV-NS3 protein. (E) Bar diagram representing the viral titers as PFU/mL in various samples. Lower panel shows the images of plaque assay plate. All the statistical analysis was carried out using the one-way ANOVA test. Data were representative of three independent experiments. *, *P* ≤ 0.05; **, *P* ≤ 0.01; ***, *P* ≤ 0.001 were considered statistically significant.

Previous report from our group suggests that TM alters AT1 and PPARγ expressions (De et al., 2022). Therefore, BHK-21 cells were pre-treated for 3 hrs as well as post-treated with GW (10µM) or AG (10µM) or TM (50µM) after infection (MOI 0.1). The cells were harvested at 24 hpi and processed for Western blot. It was observed that the JEV-NS3 level was raised by 1.5 and 2 folds in GW and AG treated samples respectively when compared to that of infection only (Fig. 6A, B, E and F). Further, AT1 protein level was increased by 1.5, 3 and 1.5, 2.5 folds in infected and treated with either GW or AG respectively as compared to mock (Fig. 6A, C, E and G). However, PPARγ level was decreased by 1, 1.5 and 2, 3.5 folds respectively in infected and treated with either GW or AG as that of mock (Fig. 6A, D, E and H). Interestingly, the levels of JEV-NS3 and AT1 proteins were decreased by 1.5 and 1fold respectively in TM treated sample as compared to infection only (Fig. 6I, J and K). Surprisingly, PPARγ expression was increased by 2-folds as compared to infection only (Fig. 6I and L).

**Fig. 6.**
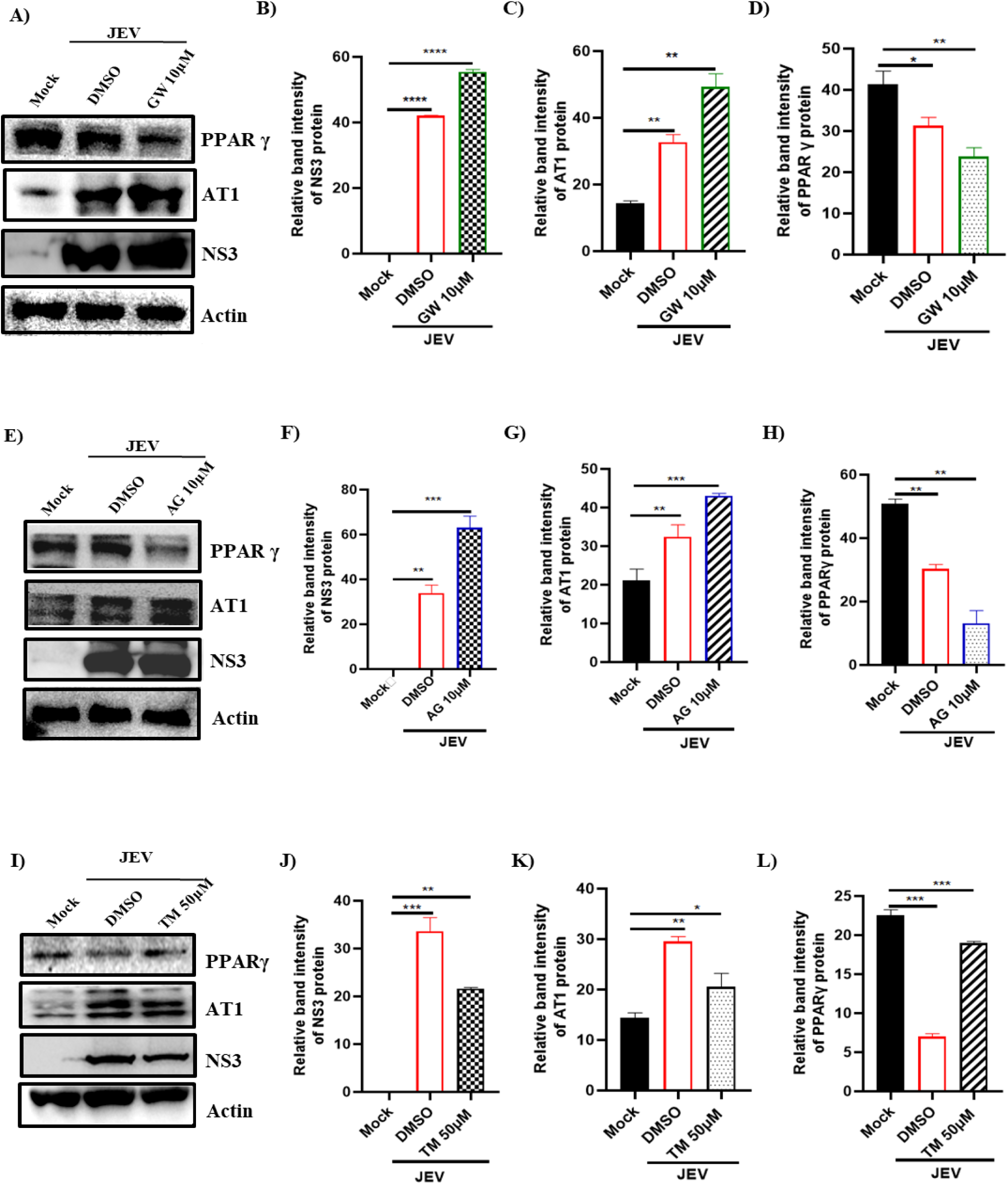
TM reduces JEV infection via AT1/PPARγ pathways: BHK-21 cells were pretreated for 3hrs with GW (10µM), AG (10µM) and TM (50µM)) separately prior to JEV infection (MOI 0.1). AG, GW and TM were added post infection also and cell pellets were harvested at 24hpi**. (**A) Western blot image depicting expressions of JEV-NS3, AT1 and PPARγ proteins in mock, infected and infected with GW treated cells. (B-D) Bar diagram depicting relative band intensities of JEV-NS3, AT1 and PPARγ proteins in different samples. E) Western blot image showing JEV-NS3, AT1 and PPARγ protein expressions in mock, infected and infected with AG treated BHK-21 cells. (F-H) Bar diagrams representing the relative band intensities of JEV-NS3, AT1 and PPARγ in different samples. (I) Western blot image representing the change in expressions of JEV-NS3, AT1 and PPARγ in mock, infected and TM treated samples. (J-L) Bar diagrams depicting relative band intensities of JEV-NS3, AT1 and PPARγ in various samples. The one-way ANOVA test analysis was performed; *, *P* ≤ 0.05; **, *P* ≤ 0.01; ***, *P* ≤ 0.001; and ****, *P* ≤ 0.0001 were considered statistically significant.

To assess the effect of the inhibitors on the cells in absence of viral infection, the BHK-21 cells were treated with TM (25 and 50µM) and cell pellet was harvested after 24 hrs. There was no significant change in the expression levels of AT1 and PPARγ when compared to the mock cells (Supplementary Fig. S3A, B and C). Similarly, the uninfected BHK-21 cells were treated with GW (10 and 20µM) and AG (10 and 20µM) for 24 hrs. Interestingly, the AT1 expression level was increased significantly in AG treated cells whereas it remained unaltered in GW treated cells as that of untreated mock control (Supplementary Fig. 4S A, B, D and E). Further, the PPARγ level remained unchanged in both AG and GW treated cells as compared to mock cells (Supplementary Fig. 4S A, C, D and F). Taken together the above findings, it can be suggested that the mechanism of action of TM against JEV is mediated through the AT1/PPARγ axis.

### 2.4. TM restricts inflammatory response induced by JEV infection via p-NF-**κ**β /COX-2/p-ERK pathways

As per Sumarno et al 2020 (Ernawati et al., 2020), neuroprotective action of TM was mediated through the involvement of p-NF-κβ /COX-2/ p-ERK (phospho-Nuclear factor kappa-light-chain-enhancer of activated B cells/ Cyclooxygenase 2/ phospho-extracellular signal-regulated kinases factors). Thus, to further understand the role of downstream players of the AT1/PPARγ pathways, JEV infection was carried out in RAW264.7 cells at MOI of 5. It was observed that expressions of GSK-3β (Glycogen synthase kinase-3beta), p-ERK1/2 and AT1 were up-regulated by 1 and 1.5 folds in infected and GW or AG treated cells respectively as that of mock (Fig. 7A, B and E). Whereas in TM treated sample, the expressions of GSK-3β, p-ERK and AT1 were down regulated significantly by 1-fold (Fig. 7A, B, C and E) when compared to infection only. The activation of p-NF-κB and COX-2, a prostaglandins maker enzyme involved in inflammation, was raised by 1 and 1.5 folds in infected and GW or AG treated and infected samples respectively as that of mock. Whereas, the p-NF-κB and COX-2 protein levels were down-regulated by 1.5 and 1 folds respectively in TM treated sample when compared to infection (Fig. 7A, F and H). The PPARγ level was reduced by 1 and 3 folds in infected and GW or AG treated samples as compared to mock. Further, in TM treated samples it was increased significantly by 2 folds as compared to infection control (Fig. 7A and G). Similarly, the JEV-NS3 protein was also reduced significantly by 2 folds when compared to infection only, whereas it was significantly upregulated in AG/GW treated RAW264.7 cells during viral infection (Fig. 7A and D). Further, there was 1-fold reduction in the level of inflammatory cytokine {p-IRF3 (phospho-interferon regulatory factor 3)} in TM treated condition as compared to infection control (Fig.7A and I). Taken together, the data indicate that TM efficiently reduces JEV induced inflammation by down regulating p-IRF-3, COX-2 and p-NF-κB.

**Fig. 7.**
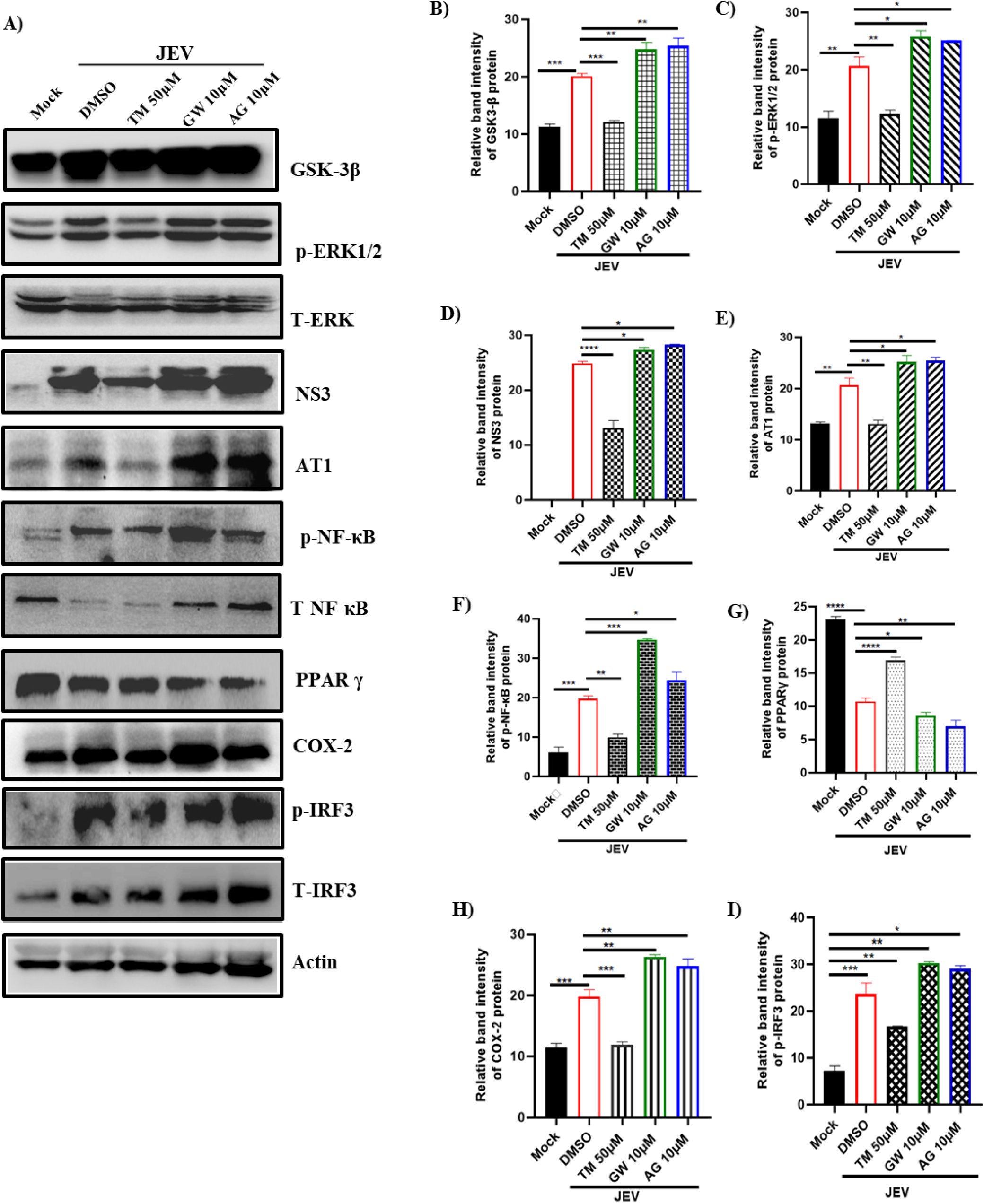
TM down-regulates inflammatory response induced by JEV infection via p-NF-κB/COX-2/p-ERK pathways: RAW 264.7 cells were pre-treated with GW (10µM), AG (10µM) and TM (50µM) for 3 hrs and then infected with JEV at a MOI of 5. The GW (10µM), AG (10µM) and TM (50µM) were again administered post-infection and cell pellets were collected after 24hpi. (A) Western blot image showing expression profiles of GSK-3β, T-ERK, NS3, AT1, T-NF-κB, PPAR γ, COX-2, T-IRF3 proteins along with phosphorylated ERK1/2, NF-κB and IRF-3 in JEV infected and TM, GW and AG treated RAW 264.7 cells. Actin was used as a loading control. (B-H) Bar diagram indicates the relative band intensity of GSK-3β, p-ERK1/2, NS3, AT1, p-NF-κB, PPAR γ, COX-2 and p-IRF3 proteins in different samples. The statistical analysis was done by the one-way ANOVA test. *, *P* ≤ 0.05; **, *P* ≤ 0.01; ***, *P* ≤ 0.001; and ****, *P* ≤ 0.0001 were considered statistically significant, n=3.

### 2.5. JEV infection was reduced by TM in mice model

Next, the anti-JEV activity of TM was evaluated in Balb/c mice (12-14 days old). The mice were infected with JEV and 10mg/Kg dose of TM was administered orally at an interval of 24 hrs upto 6 dpi (Fig. 8A). The JEV infected mice showed rise in body temperature, ruffled fur, hunch back, slow movement and hind limb paralysis, whereas TM treated animals showed no such symptoms (Fig. 8B). Further, expression of the JEV-NS3 protein was reduced drastically (>50%) in TM treated mice brain as that of infected one (Fig. 8C and D). The clinical score graph showed lessening of symptoms in TM treated mice (Fig. 8E) and immunohistochemistry analysis revealed decrease in the JEV-NS3 protein level in drug treated mice brain when compared to infection only (Fig. 8F). Interestingly, JEV infected brain tissue section showed increased number of perivascular cuffs and microglia nodules which were found to be reduced significantly in TM treated section (Fig. 8G). Thus, the above findings indicate that TM can reduce JEV infection and protect mice efficiently.

**Fig. 8.**
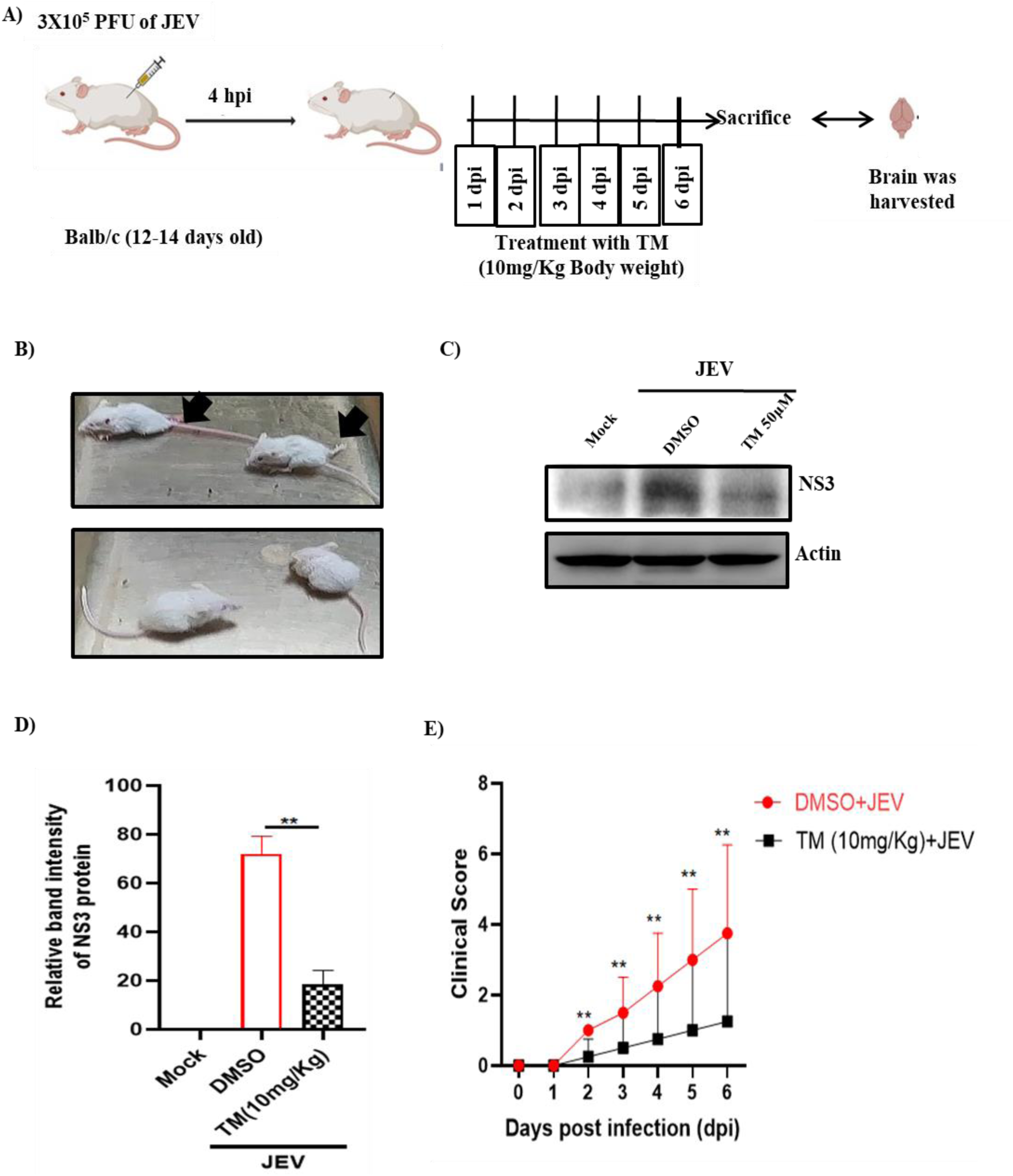
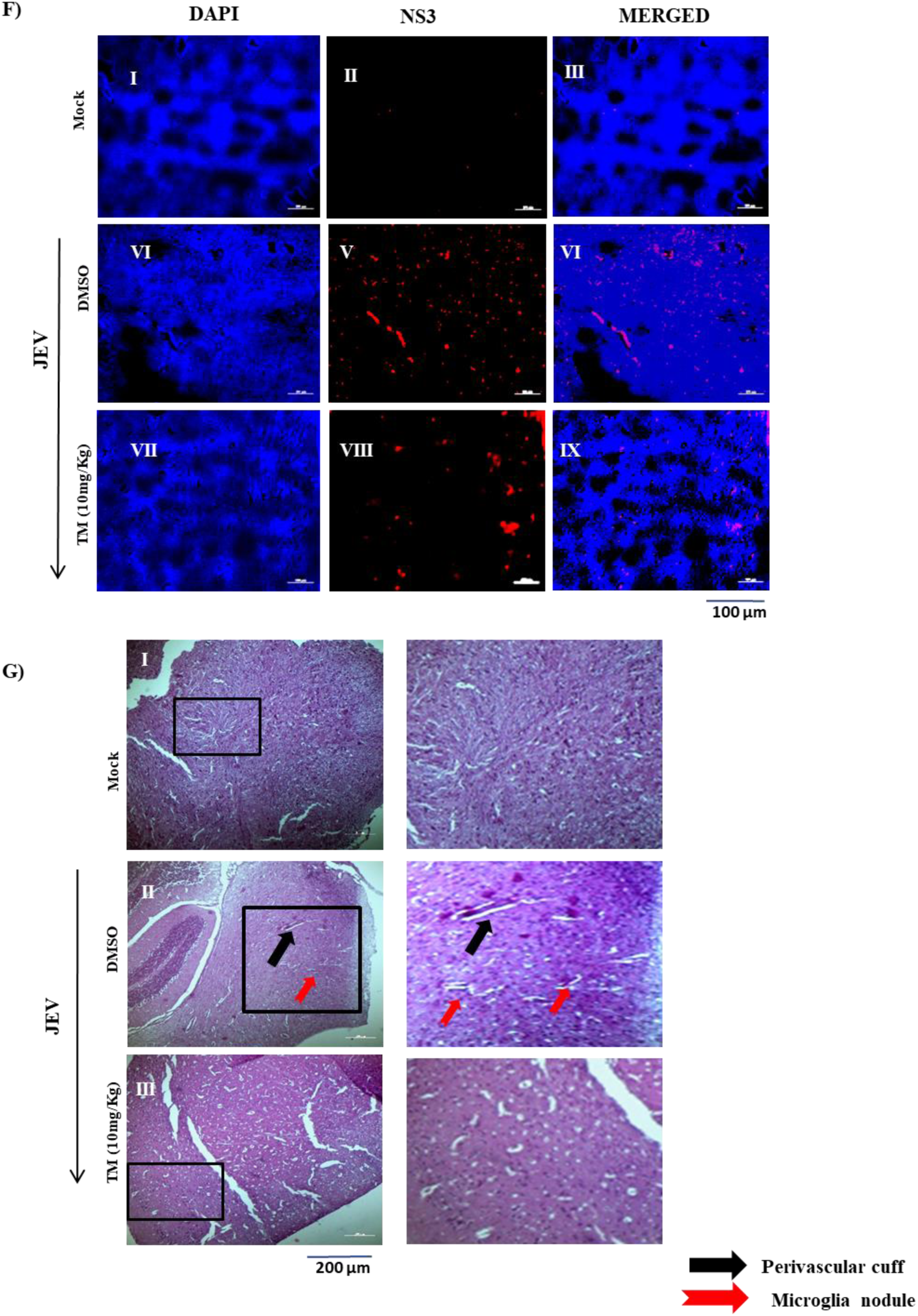
TM reduces JEV infection in mice model: The Balb/c (12-14 days old) mice were injected subcutaneously with 3X10^5^ PFU of JEV and TM (10mg/Kg) was given orally at an interval of 24hrs upto 6 days. Mice were sacrificed on 7^th^ day post infection; sera and brain tissue were collected for further downstream processing. (A) Schematic diagram representing experimental workflow in mice model. (B) Image representing JEV infected and drug treated mice and arrow indicating hind limb paralysis in infected mice at 6^th^ dpi. (C) Western blot image depicting the JEV-NS3 protein in brain tissue sample of mock, infected and treated mice. (D) Bar diagram indicating relative band intensity of JEV-NS3 protein in different samples. (E) Graph showing the clinical scores of the disease symptoms of JEV infected and treated mice which were under observation from 1 dpi to 6 dpi (*n* = 4). (F) Confocal images representing mock, JEV infected and infected along with treated brain tissue samples stained for the JEV-NS3 protein (Scale Bar 100µM). (G) Image showing H&E-stained mock, infected and drug treated brain tissue samples, black arrow indicating perivascular cuffs and red arrow indicating microglia nodule. Scale bar 200 µM. Data of three independent experiments is shown as mean ± SD.

## 3. Discussion

In the Asia-Western Pacific region, JEV is the most prominent cause of encephalitis, which is associated with high mortality or neuropsychiatric sequelae (Joe et al., 2022). While the vaccines have failed to eradicate JEV, efforts have not been successful in developing a clinically effective antiviral strategy. Considering the proinflammatory nature of the AT1 axis in inducing encephalitis, ARBs can help managing JEV infection. Because of the relatively higher ability to cross BBB, TM can be prioritized amongst the ARBs. It is also a partial agonist of PPARγ, which has been reported to be downregulated during viral encephalitis (Layrolle et al., 2021). Other than mediating encephalitis, the AT1/ PPARγ axes are also critical host factors for the replication of several alphaviruses and flaviviruses. Considering TM’s ability to modulate these axes, there is great interest in investigating its efficacy against JEV infection.

In the current study, TM was found to be non-cytotoxic to BHK-21 cells till 160µM and efficiently inhibited JEV infection (MOI of 0.1) in the BHK-21 cell line with 50 µM (90%) and 75µM (95%) concentrations following post-infection. This was also associated with a significant reduction in viral RNA (50% at 50 µM; 75% at 90 µM) and protein levels (52% at 50 µM; 75% at 90 µM). This anti-JEV property was also supported by reduced expressions of JEV-E, iNOS, and Cas3 genes in physiologically relevant cell lines, including macrophages (RAW264.7) and neuronal (SH-SY5Y) cells. Interestingly, TM was able to reduce the viral load in pre, co, and post-treatment conditions. However, the reduction was more (80%) in post-treatment conditions. Interestingly, viral titer was also reduced significantly (30%) when the virus particles were incubated with TM. Thus, some virus inactivation activity of TM cannot be ruled out. TM is known to be an AT1 blocker and partial PPARγ agonist. So, to understand the involvement of these axes in mediating TM’s antiviral action against JEV, agonists of AT1R (AG) and antagonists of PPARγ (GW) were used. Viral titer was enhanced (80% and 90%) in presence of GW/AG, suggesting that TM could abrogate viral infection through the AT1/PPARγ axis. The AT1 protein level was upregulated (1.5, 3, and 2.5 folds), whereas the PPARγ level was downregulated (2, 1.5, and 3.5 folds) during infection and with GW or AG treated conditions. Interestingly, the AT1 (1-fold) and PPARγ (2 folds) levels were reversed in presence of TM. Additionally, TM showed a significant reduction in the inflammatory responses by downregulating p-IRF-3, COX-2, and p-NF-κB. Further evaluation in Balb/c mice model revealed that TM can remarkably reduce the disease score, viral protein level and histological changes in brain tissue of JEV infected mice.

TM showed a dose-dependent reduction in viral titer with about 95% abrogation of virus particle formation with 75µM concentration when administered post-JEV-infection (0.1 MOI). A proportionate decrease in viral RNA and JEV-NS3 protein levels was also observed. Further, CC_50_ of TM was found to be > 350μM. A similar CC_50_ has been reported for TM in Vero and RAW264.7 cells (De et al., 2022) . With an IC_50_ of 24.68μM, the selectivity index of TM was estimated to be > 14.18 (Fig. 2). This selectivity index and the fact that the clinical safety profile of TM is well-established suggests its potential for repurposing. In further support of this, the effect of TM was demonstrated in physiologically relevant cell lines, including macrophages (RAW264.7) and neuronal (SH-SY5Y) cells. A significant reduction was observed in the expressions of JEV-E in a dose-dependent manner. Moreover, TM showed substantial reductions in INF-β gene expression and iNOS levels in RAW 264.7 cells. Since INF-β (Barrett et al., 2020) and iNOS (Liy et al., 2021) are markers of neuroinflammation, this also indicates the neuroprotective potential of TM, which is necessary for the management of JE (Şen & Hacıosmanoğlu, 2022). Also, Cas3 genes were significantly reduced by TM in neuronal cell lines (SH-SY5Y), indicating decreased apoptosis. Taken together, these findings suggest potential of TM to manage JE.

Although TM reduced the viral load in pre- and co-treatment conditions, the effect was more pronounced in post-treatment conditions with >80% reduction in viral titer. Higher efficacy in post-treatment indicates some host-mediated action. Nonetheless, it also showed a 30% abrogation of virus particles following incubation with TM, which suggests some direct antiviral effects. However, no specific virus targets can be identified at this point. TM has earlier been reported to directly affect the protease activity for CHIKV-nsP2 (Tripathi et al., 2020). However, the same cannot be said about JEV and this needs further investigation. Hence, TM can be said to have predominately host-mediated antiviral properties against JEV. Nevertheless, considering its significant reduction in pre-treatment conditions, the involvement of some viral targets cannot be ruled out.

To understand the mechanisms of action of TM, the primary mediator of its pharmacodynamics, AT1 protein and PPARγ were monitored. The AT1 protein expression was enhanced in infected cells, whereas PPARγ expressions were downregulated. This suggests the involvement of both these axes in JEV infection. Further, AT1 agonist (AG) and PPARγ antagonist (GW) augmented the viral infection. Entry and replication of Flaviviruses are reported to be modulated by AT1 activation (Pedreañez et al., 2024). For example, C-type lectin on dendritic cells called dendritic cell intercellular adhesion molecule 3-grabbing nonintegrin (DC-SIGN) and glucose-regulated protein-78 (GRP78), which play a critical role in JEV entry as well as replication (Anwar et al., 2022; Nain et al., 2017) are reported to be upregulated with AT1 activation (Pedreañez et al., 2024) during infection of JEV as well as some other Flaviviruses including DENV. Hence, the upregulation of the AT1 protein may have contributed to the entry and replication of JEV infection. Although there is no specific data about the involvement of PPARγ in JEV entry or replication, it has pleiotropic roles in viral infections and inflammations in the brain (Layrolle et al., 2021). Thus, the reduction in PPARγ expression may have contributed to the rise in infection in multiple ways, which need further investigation. In the present study, the AT1 level was downregulated, and the PPARγ level was upregulated upon TM treatment. Altogether, the above findings suggest the pivotal role of AT1 and PPARγ in the inhibitory action of TM against JEV, which can be further explored for therapeutic intervention for multiple viral infections.

To further support this mode of action, some downstream players of AT1 and PPARγ were explored. Key downstream players of AT1/PPARγ pathway were analyzed, demonstrating the reduction of inflammatory markers such as p-IRF3, GSK-3β, p-ERK1/2, p-NF-κB and COX-2. The gene expression levels of GSK-3β, COX-2, and p-NF-κB proteins are reported to be higher in JEV and their reductions has been associated with decrease in JEV-infection in human microglia (CHME3) and human neuroblastoma cells (SH-SY5Y)) (Kumar et al., 2020). Thus, it can be proposed that in JEV infection, AT1 was upregulated, which in turn activates GSK-3β. At the same time, viral infection reduces PPARγ, which triggers the increased production of transcription factors such as p-IRF3, p-NF-κB, and pro-inflammatory markers COX-2. Finally, it results in increased viral infection, cell injury, and death (Fig. 9A). On the other hand, the presence of TM leads to a reduction of AT1, pro-inflammatory cytokines, GSK-3β, an increase in PPARγ resulting in abrogation of viral infection and inflammation (Fig. 9B).

**Fig. 9.**
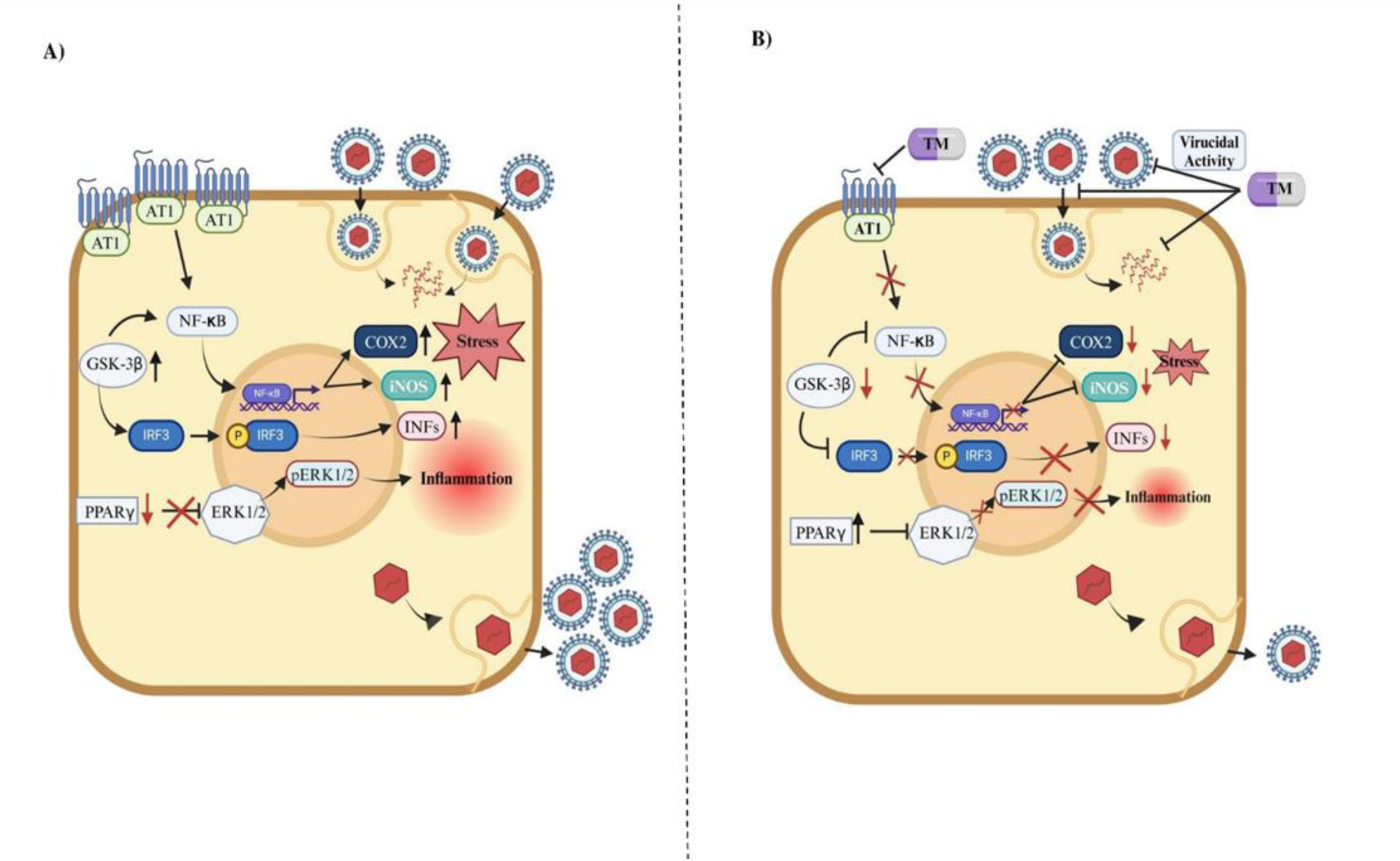
Diagrammatic representation of proposed working model of TM: RAW264.7 cells were infected with JEV (MOI 5) and 24 hpi cells were harvested. **(A)** Image showing JEV infection alters expressions of AT1 (increased), PPARγ (reduced) and other downstream key players such as p-ERK1/2, p-NF-κB, p-IRF3, GSK-3β and COX-2. (B) Image representing TM treatment leads to decreased AT1 and increased PPARγ expressions which further leads to decreased phosphorylation of ERK1/2, NF-κB and IRF3 and also down regulates the expression of GSK-3β as well as COX-2. Also, TM interferes with JEV life cycle, entry and also possesses virucidal activity results in decreased viral load. This explains reduction in oxidative stress and inflammation preventing cell injury and death.

Further, it has been reported by Rong Jiang, Jing Ye et al 2014 that p-ERK1/2 level is upregulated in JEV infection. Moreover, JEV infection leads to the overproduction of ROS and activation of nuclear factor kappa B (NF-κB) and other signaling pathways such as (Apoptosis signal**-**regulating kinase 1) ASK1-ERK/p38 MAPK (Mitogen-activated protein kinase) signaling pathways (Zhang et al., 2014) leading to inflammation. Also, the AT1 axis mediates inflammation through NF-κB/cytokine pathway (Ang et al., 2012). Besides, this host mediated antiviral effects, TM directly impedes JEV infectivity. Taken together, TM inhibits viral infection directly and indirectly by modulating host factors and managing the JEV-induced inflammation.

The *in vitro* intervention was co-related with *in vivo* studies. With once-a-day oral administration of TM (10mg/Kg) for six days, the JEV NS3 protein expression was reduced drastically (50% approximately) in the brain of the Balb/c mice, suggesting a significant reduction in JEV infection of the brain. The reduction in encephalitis was evident from a significant decrease in perivascular cuffs and microglia nodules. Since encephalitis and neurological sequelae are the primary reasons for mortality and morbidity, these findings demonstrate the efficacy of TM in the management of JE. Previous reports suggest the high dose of drugs NTZ (50 mg/kg/day, 75 mg/kg/day and 100 mg/kg/day), and AV1004 (35 mg/kg) for 25 and 23 days to decrease JEV load respectively (Shi et al., 2014; Zhang et al., 2021). Similarly, a report demonstrated that 15mg/kg twice daily berberine dose was given to 3-4 week old BALB/c mice for up to 15 days to reduce JEV infection (Huang et al., 2021). Compared to these, TM was effective even after a shorter duration (6 days) of treatment. TM is widely used for prolonged therapy of hypertension at a dose of 40mg/day or higher. Thus, the effectiveness of TM at the equivalent of this human dose in mice (10mg/kg) suggests the possibility of its acceptability while repurposing. This is also helpful because, at these lower doses, TM is relatively safe in normotensive individuals (Makino et al., 2008). However, for effective repurposing, the dose regimen needs further optimization. The oral bioavailability of TM is dose-dependent (40% at 40 mg/day and 58% at 160 mg/day) (Paramanindita et al., 2020). Since increasing doses for normotensive JEV patients may have adverse impacts, it is necessary to improve bioavailability at lower doses for repurposing application. Also, morbidity and mortality of JEV are primarily higher in children and young adults, hence the safety profile of TM in these groups needs to be evaluated. Alternatively, selective delivery of TM to the brain needs to be designed for effective repurposing against JEV.

## 4. Conclusion

Both in vitro and in vivo findings show the ability of TM to reduce JEV infection. The AT1 and PPARγ axes, known to mediate encephalitis, were also found to be involved in JEV infection. Besides modulating these axes, TM may also interfere with some antiviral targets for its anti-JEV properties. Further, TM can potentially manage JE by reducing the downstream markers of these axes that mediate inflammation. The preclinical efficacy at a human equivalent dose and with a relatively shorter duration of exposure suggests its suitability for further repurposing.

## 5. Materials and Methods

### 5.1. Cells and Virus

The BHK-21 and RAW264.7 cells were procured from the National Centre for Cell Sciences (NCCS); Pune, India whereas the PS (Porcine Stable Kidney) and SH-SY5Y cell lines were kind gift from Dr A. Basu, (NBRC; Gurgaon, India) and Dr G. H. Syed (Institute of Life Sciences, Odisha) respectively. The BHK-21 and PS cells were maintained in Dulbeco’s modified Eagle’s medium (DMEM; Himedia, India) and RAW264.7 cells were maintained in Roswell Park Memorial Institute media (RPMI 1640, Gibco Gluta MAX; Invitrogen, Cal, US) respectively along with 10% fetal bovine serum (Gibco, Invitrogen), gentamycin and penicillin-streptomycin (Sigma, USA). The SH-SY5Y (neuronal cells) was maintained in DMEM/F-12 media (Gibco™). All the cells were kept in 5% CO_2_ incubator at 37°C. The GP78 strain (accession No. **<u>AF075723</u>**) of JEV was a kind gift from Dr A. Basu (NBRC; Gurgaon, India).

### 5.2. Antibodies and compounds

The JEV-NS3 (Non structural protein-3) and actin antibodies were obtained from GeneTex (Cat. No. GTX125868) and Sigma-Aldrich (MO, US) respectively. The PPARγ (Peroxisome proliferator-activator receptor-gamma), T-ERK1/2 (total-extracellular signal-regulated kinase), p-ERK1/2 (phospho-extracellular signal-regulated kinase), p-NF-κB, T-NF-κB (total-Nuclear factor kappa-light-chain-enhancer of activated B cells), COX-2, GSK-3β (glycogen synthase kinase-3 beta), p-IRF3, T-IRF3 (total-interferon regulatory factor 3), were procured from Cell Signaling Technology (CST) (MA, US). The AT1 antibody was obtained from Santa Cruz Biotech (TX, US). The [Val5]-angiotensin II acetate salt hydrate (AG) is an ANG II analog having valine instead of isoleucine in an ANG II octapeptide at fifth amino acid position (Sigma-Aldrich, MO, US) and GW9662 (GW) is an irreversible PPARγ antagonist (Sigma-Aldrich, MO, US). Telmisartan (TM) was a kind gift from Glenmark Life Sciences Ltd., Ankleshwar, Gujrat, India.

### 5.3. Virus Infection

Cells were seeded onto plate with 70-80% confluency. The JEV infection was carried out at 0.1 MOI in BHK-21 and SH-SY5Y cells, whereas at Multiplicity of Infection (MOI) of 5 in RAW264.7 cells in serum free media (SFM) with shaking at an interval of every 15 mins up to 90 mins as per (Mishra et al., 2016). The cells were washed twice with 1X PBS and complete media was then added along with the TM at varying concentrations post infection. The cells were incubated at 37 °C with 5% CO_2_ till 24 hours post infection (hpi). The cell culture supernatant and pellets were harvested at 24hpi for further downstream processing.

### 5.4. Plaque assay

The plaque assay was performed as described earlier by (De et al., 2022). Briefly, the cell supernatant from mock, infected and treated samples were collected and serially diluted in SFM. The diluted samples were added onto monolayer of PS cells and infection was carried out as described before. After infection, the cells were overlaid with carboxymethyl cellulose (Sigma-Aldrich, MO, US). The plates were incubated at 37°C with 5% CO_2_ for 6-7 days until clear plaques were visible. The cells were then fixed with 8% formaldehyde (Sigma-Aldrich, St. Louis, MO) and stained with 0.1% crystal violet (Sigma-Aldrich, St. Louis, MO). The number of plaques were calculated and presented as plaque-forming units per milliliter (PFU/mL).

### 5.5. Cytotoxicity assay

The MTT assay was carried out using the EZcountTM MTT cell assay kit (Himedia, Mumbai, India) as per manufacturer’s protocol. Approximately, 10,000 BHK-21 cells were seeded in 96 well plates 24 hrs prior to infection. The cells were then treated with different concentrations of TM (5, 10, 20, 40, 80, 160, 250 and 350µM) or GW (1, 5, 10 and 20 µM) or AG (10, 20, 30 and 40µM) for 24 hours along with a reagent control. The absorbance was taken at 570 nm. The percentage of metabolically active cells was compared with the control cells to determine cellular cytotoxicity 50 (CC_50_) as mentioned previously by (De et al., 2022).

### 5.6. Western blot

The Western blot analysis was done as described previously by (Chatterjee et al., 2022). In short, the mock, infected and treated samples were harvested and lysed with RIPA lysis buffer. Similarly, mock, infected and treated brain tissue was snap-frozen and lysed by the homogeniser in RIPA buffer. Equivalent concentration of each sample was loaded and run on 10% SDS-polyacrylamide gel. The proteins from cell culture supernatant and brain tissue samples were transferred onto polyvinylidene difluoride (PVDF) membrane and probed with primary antibodies JEV-NS3, T-ERK1/2, p-ERK1/2, p-NF-κB, T-NF-κB, COX-2, GSK-3β, p-IRF3, T-IRF3 in 1:1000 dilutions, AT1 (1:250) and PPARγ (1:250) for overnight. Further, brain tissue sample was probed with primary antibodies JEV NS3 (1:1000) and AT1 (1:250) overnight. Actin (1:500) was used as an internal control. The detection was done by corresponding secondary antibodies tagged with horseradish peroxidase (HRP). The blots were imaged by Chemidoc (BioRad) and protein band was quantified using the Image J software (n=3).

### 5.7. Confocal microscopy

The immunofluroscence assay was performed as described previously by (Chatterjee et al., 2022). The BHK-21 cells were seeded onto coverslip in 6 well plate with 50% confluency. The cells were infected as described above. At 24hpi, the cells were fixed with 4% paraformaldehyde and incubated with primary antibody JEV-NS3 (1:2000) for 45 mins. After washing the cells with 1XPBS, secondary antibody 594 anti-rabbit Alexa Flour (Invitrogen) were added for 45 mins. The cells were mounted with antifade reagent (Invitrogen) to avoid photobleaching. The images were acquired using the Leica TCS SP5 confocal microscope (Leica Microsystems, Heidelberg, Germany) with a 10X objective.

### 5.8. Treatment of TM pre infection, co infection and post infection

The pre, during and post assay was performed as described earlier by (De et al., 2022). In brief, BHK-21 cells were pretreated with TM (50µM) for 3hrs and later on infection was carried out. In during condition, the drug was given along with the virus and then infection was carried out for 1.5 hrs. Further, in the post treatment, drug was added after the virus infection. The supernatant from all the above mentioned three conditions were collected at 24hpi and viral titre was determined by plaque assay.

### 5.9. Virucidal Assay

The virucidal activity of TM was determined as mentioned previously by (Kitidee et al., 2023). In brief, virus (0.1 MOI) was incubated with 50 µM TM for 1 hr in serum free media at 37°C in 5% CO_2_ incubator. The plaque assay was performed with the untreated and drug treated inoculums to determine the viral titer.

### 5.10 Treatment with compounds (GW or AG)

The BHK-21 cells were pre-treated for 3 hrs with increasing concentrations of GW (1, 5, 10 and 20 µM) or AG (10, 20, 30 and 40 µM) as described previously by (De et al., 2022). Later on, infection with JEV (MOI 0.1) was given as mentioned above. Post infection, again GW or AG was administered with the above stated concentrations and samples were collected at 24 hpi.

### 5.11. Flow cytometry

The flow cytometry analysis was done as described earlier by (Sanjai Kumar et al., 2021). In brief, mock, infected and infected along with TM-treated cells were harvested at 24 hpi. These cells were fixed with 4%paraformaldehyde followed by resuspension in FACS buffer (1X PBS, 1% BSA, 0.01% NaN_3_ (Sigma-Aldrich, St. Louis, MO). The cells were treated with permeabilization buffer (1X PBS, 0.5% BSA, 0.1% Saponin and 0.01% NaN_3_) followed by blocking with 1% BSA at room temperature for 30 mins. The primary antibody anti-rabbit JEV NS3 (1:1000) was added for 45 mins. Next, the cells were washed with permeabilization buffer, followed by addition of fluorescent labelled secondary Alexa Fluor (AF) 594-conjugated anti-rabbit antibody (Invitrogen, US antibody). Approximately, 10,000 cells were acquired by the LSR-Fortessa Flow cytometer (BD Biosciences, CA, US) for each sample and analysed by the FlowJoV10.7.1. (BD Biosciences, CA, US).

### 5.12. qRT-PCR

The viral and total RNA was extracted from supernatant and cells using the QIAamp viral RNA mini kit (Qiagen, Hilden, Germany) and TRIzol (Sigma) respectively as described earlier by (Chatterjee et al., 2023). The cDNA (PrimeScript 1^st^ strand cDNA synthesis kit, Takara, Japan) was synthesized using equal volume of viral RNA and 1µg of total RNA as per manufacturer’s instructions. The qRT-PCR was performed using the JEV-E (envelope) (Datey et al., 2020), interferon-β (INF-β), iNOS (Inducible nitric oxide synthase), (Wang et al., 2019) and Cas3 (Caspase3) (Kitidee et al., 2023) genes specific primers (Supplementary Table No.2) using Mesagreen Sybr green master mix (Eurogentec) as per manufacturer’s protocol. GAPDH was used as control (Supplementary Table No.1). The copy number per mL was calculated as described previously by (Sanjai Kumar et al., 2021) Sanjai et al whereas the change in gene expression was calculated by the 2^−ΔΔ CT^ method.

### 5.13. Animal Studies

All the animal work was carried out under the strict guidance of Committee for the Purpose of Control and Supervision of Experiments on Animals of India. Every procedures and tests were evaluated and permitted by the Institutional Animal Ethics Committee (Reference No. ILS/IAEC-320-AH/MAY-23).The JEV infection was performed in 12-14 days old Balb/c mice as described previously by (Swarup et al., 2007) (doi:10.1128/AAC.00041-07). In short, a lethal dose of 3×10^5^ PFU of JEV (GP78 strain) was injected subcutaneously. After 4 hours post infection, drug was given orally at a dose of 10mg/Kg at an interval of 24 hours upto 6 days post infection (dpi) to the treated mice group (n=4). At 7^th^ dpi the mice were sacrificed, sera were separated from blood and subjected to qRT-PCR. Brain tissue and serum were collected and processed for Western blotting and qRT-PCR respectively. The clinical score was recorded for each mouse depending upon the onset of symptoms on daily basis (0, no symptoms; 1, ruffled fur; 2, hunch back; 3, slow movement; 4, hind limb paralysis; 5, death).

### 5.14. Histopathological analysis

was carried out as described earlier by (De et al., 2022) with little modification in the protocol. For histopathological examinations, formalin-fixed tissue samples were dehydrated and embedded in paraffin wax, and serial paraffin sections of 5mM were obtained. The cut sections were then heated at 54°C in water bath and then dried at 37°C overnight (Robertson et al., 2008). The sections were then stained with hematoxylin and eosin (H&E) (Himedia Cat no. S014-500ML and S007-500ML respectively), and histopathological changes were visualized using a light microscope (Zeiss Vert.A1, Germany). Sections were also examined for the presence of JEV-NS3 protein using a specific antibody. Briefly, NS3 antibody (1:750) was added to the slides overnight at 4°C. After washing with 0.1% PBST thrice, slides were incubated with Alexa Fluor 594 (1:750; anti-rabbit; Invitrogen, MA, US) for 45 mins -1 hr at room temperature in the dark and humidified chamber. The slides were washed thrice with 0.1% PBST and then mounted with a mounting reagent using DAPI (Invitrogen, MA, US).

### 5.15. Statistical Analysis

The statistical analyses were performed using the GraphPad Prism version 8.0.1 software. Data of three independent experiments are shown as mean ± SD.*P <0.05, **P <0.01, *** P <0.001, ***** P <0.0001 were considered as statistically significant. For three groups one way analysis of variance (ANOVA) with Brown-Forsythe test was used whereas for comparing two values Student’s *t* Test was carried out. All the experiments were performed in triplicates.

## CRediT authorship contribution statement

**Ankita Datey:** Conceptualization, investigation, methodology, writing original draft, data curation, validation; **Sanchari Chatterjee:** Investigation and data curation; **Soumyajit Ghosh**: Investigation, methodology, writing original draft and data curation; **P Sanjai Kumar:** Investigation and data curation; **Saikat De:** Investigation and conceptualization; **Udvas Ghorai:** Investigation and data curation; **Bharat Bhusan Subudhi:** Conceptualization, writing, review and editing, supervision; **Soma Chattopadhyay:** Conceptualization, writing, review and editing, supervision.

## Acknowledgement

We are thankful to Dr Anirban Basu and Dr Gulam Hussain Syed for providing the JEV (GP78) virus strain and SH-SY5Y cell line respectively. Ankita Datey and Soumyajit Ghosh are supported by CSIR-DIRECT SRF (File No. 09/0657(18482)/2024-EMR-I) and ICMR-SRF (Letter No. VIR/Fellowship/2/2022-ECD-I) fellowships, Government of India respectively. We thank ILS core grant for financial support. We would like to acknowledge Dr Punit Prasad, ILS Bhubaneswar for sharing the Biorender license for generating working model. We also thank Rajshree Rajmohan Jena for helping in the animal experiments.

## Data availability

The data that support the findings of this study are available from the corresponding author upon reasonable request. Some data may not be made available because of privacy or ethical restrictions.

---

**Fig. 1S.**
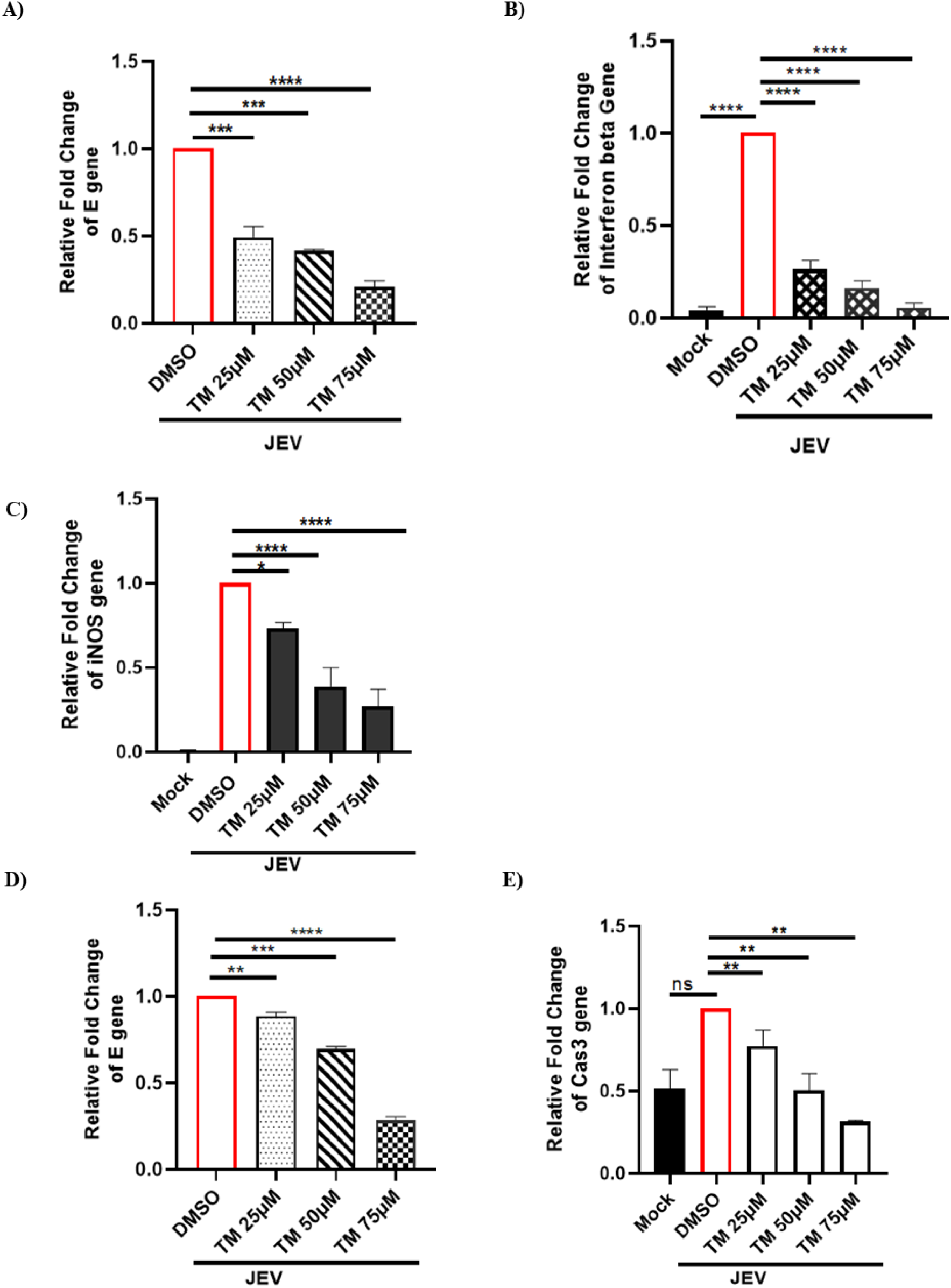
TM efficiently reduces JEV load in physiologically relevant cell lines: The RAW264.7 and SH-SY5Y cells were infected with JEV at a MOI of 5 and 0.1 respectively. After 24hpi cell pellets were harvested in TRIzol and total RNA was extracted. The qRT-PCR was performed with JEV-E (envelope) and Interferon beta (INF-β) genes specific primers. (A) Bar diagram representing the relative fold change of E gene in RAW264.7 cells (mock, infected and treated with different concentrations of TM). (B and C) Bar diagrams showing the relative fold change in INF-β and iNOS levels in mock, infected and treated RAW264.7 cells with varying concentrations of TM. (D and E) Bar diagram depicting reduction E and Cas3 gene expression in treated samples as compared to infection only in SH-SY5Y cells. The one-way ANOVA test was performed for statistical analysis *, *P* ≤ 0.05; **, *P* ≤ 0.01; ***, *P* ≤ 0.001; and ****, *P* ≤ 0.0001 were considered statistically significant, n=3.

**Fig. 2S.**
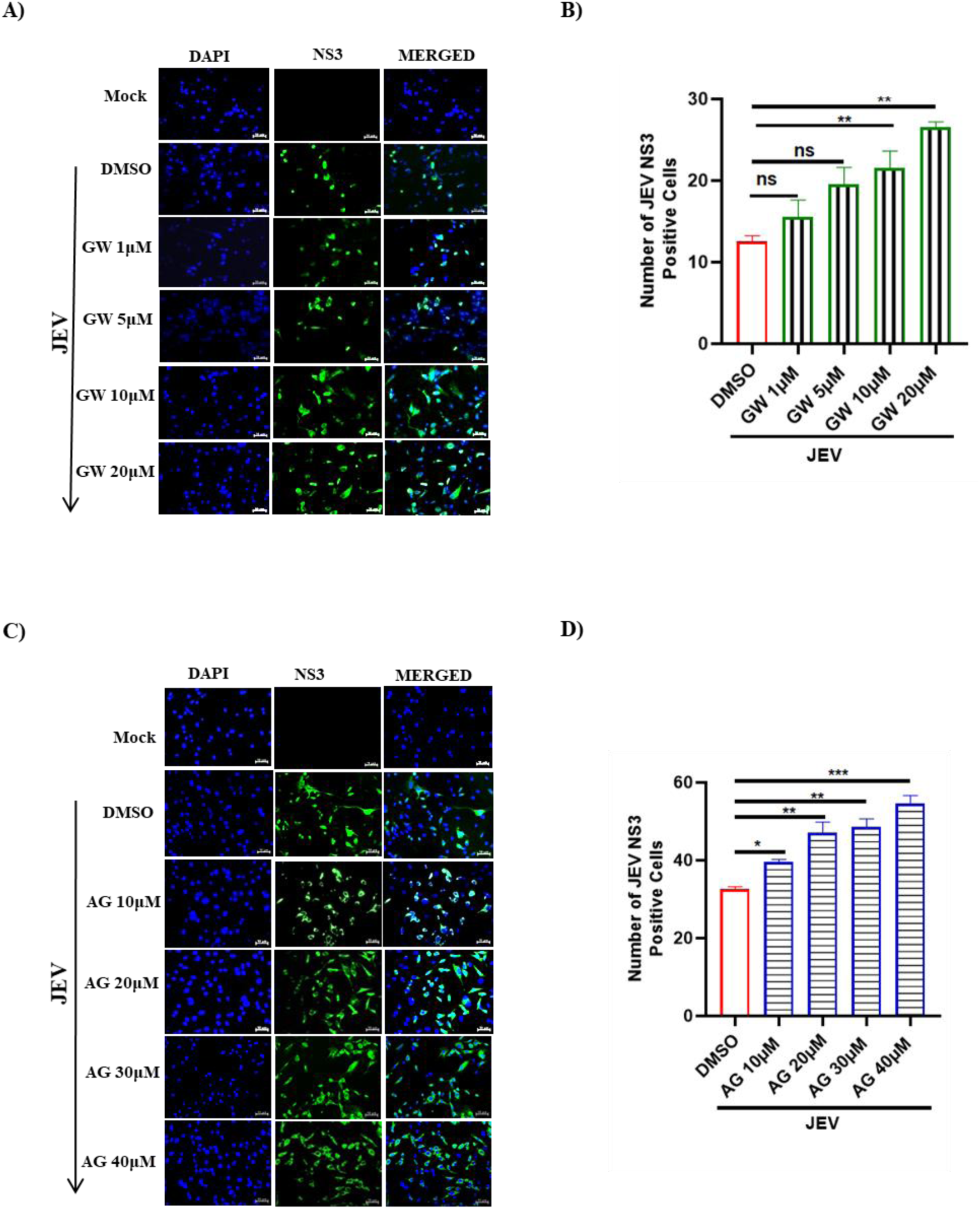
Viral load abrogates with increasing concentrations of GW or AG: BHK-21 cells were seeded onto coverslip and GW/AG were added pre and post infection. The infection was carried out at 0.1 MOI. The cells were fixed after 24hpi and incubated with JEV-NS3 (1:2000) primary antibody. (A and C) Confocal images showing number of JEV infected cells with increasing concentration of GW and AG respectively. (B and D) Bar diagram representing number of JEV-NS3 positive cells. The statistical analysis was done using the one-way ANNOVA. *, *P* ≤ 0.05; **, *P* ≤ 0.01; ***, *P* ≤ 0.001; were considered statistically significant whereas ns denotes non-significant.

**Fig. 3S.**
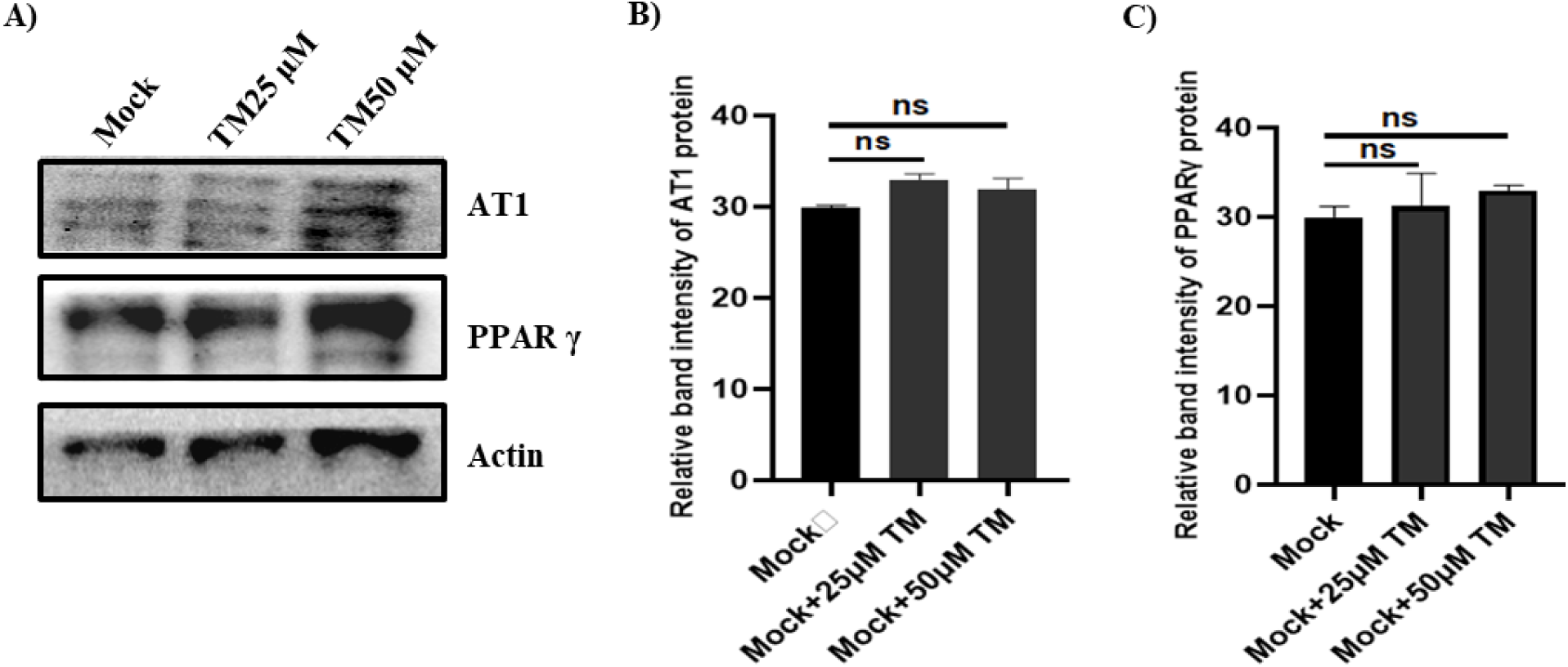
AT1 and PPARγ expressions remain unchanged after TM treatment in mock cells: BHK-21 cells were treated with TM (25 and 50µM) and after 24hpi cells were harvested, lysed with RIPA buffer. (A) Western blot image depicting expressions of AT1 and PPARγ proteins in TM treated mock cells. (B-C) Bar diagram showing relative band intensities of AT1 and PPARγ proteins. Actin was used as a loading control. The statistical analysis was done by the one-way ANOVA test, (ns=non-significant).

**Fig. 4S.**
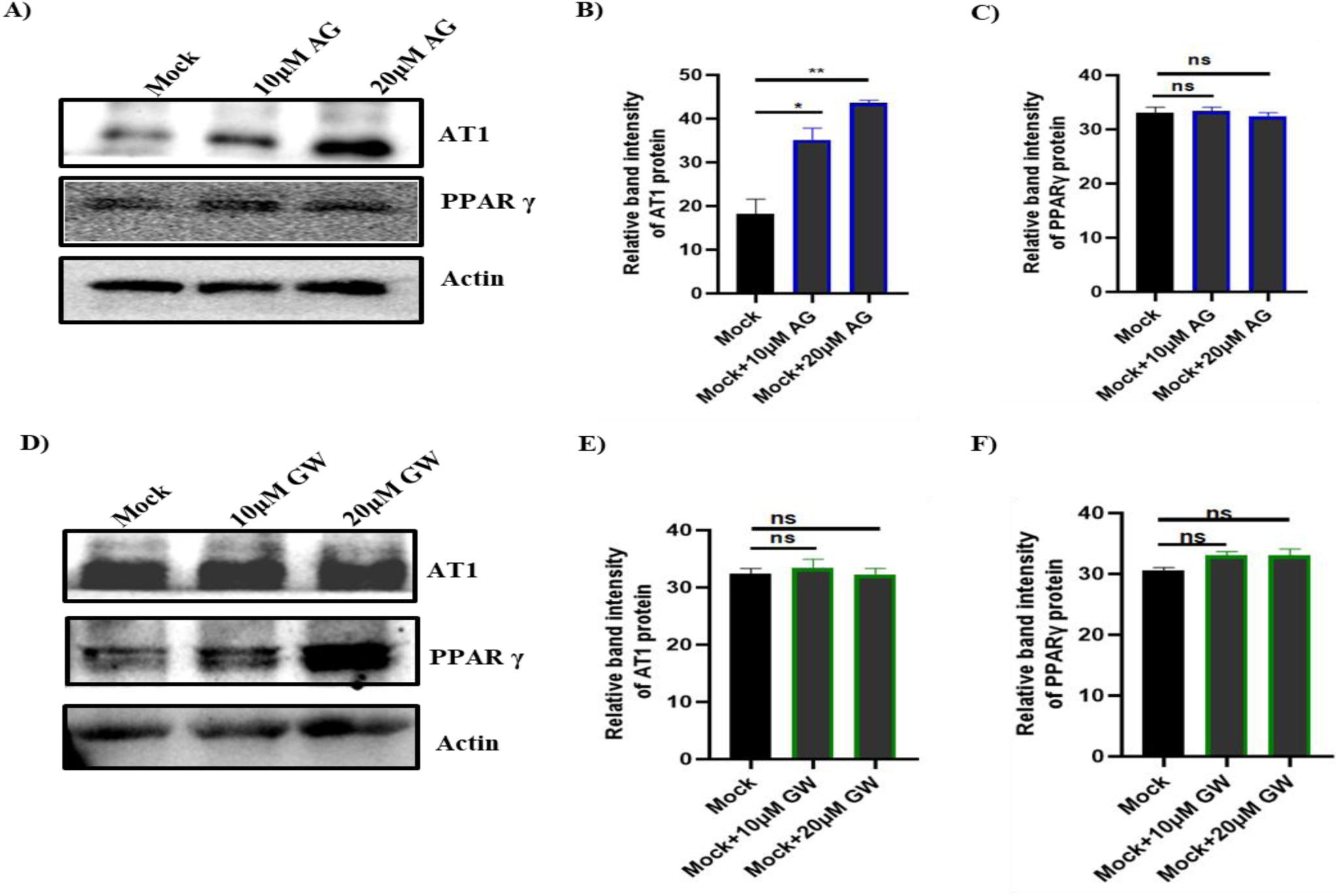
Expression profile of AT1 and PPARγ proteins in presence of AG and GW inhibitors: BHK-21 cells were incubated with AG and GW inhibitors (10 and 20µM) and cell pellets were collected after 24 hpi. (A and D) Western blot image representing AT1 and PPARγ expression in AG and GW treated cells respectively. (B and E) Bar diagram showing relative band intensities of AT1 protein in AG and GW treated mock cells respectively. (C and F) Bar diagram indicating relative band intensities of PPARγ protein in AG and GW treated BHK-21 mock cells respectively. Actin was used as a loading control. The analysis was carried out using the graphpad prism tool by the one-way ANOVA test. *, *P* ≤ 0.05; **, *P* ≤ 0.01; were considered statistically significant, n=3.

**Supplementary Table No.1:**
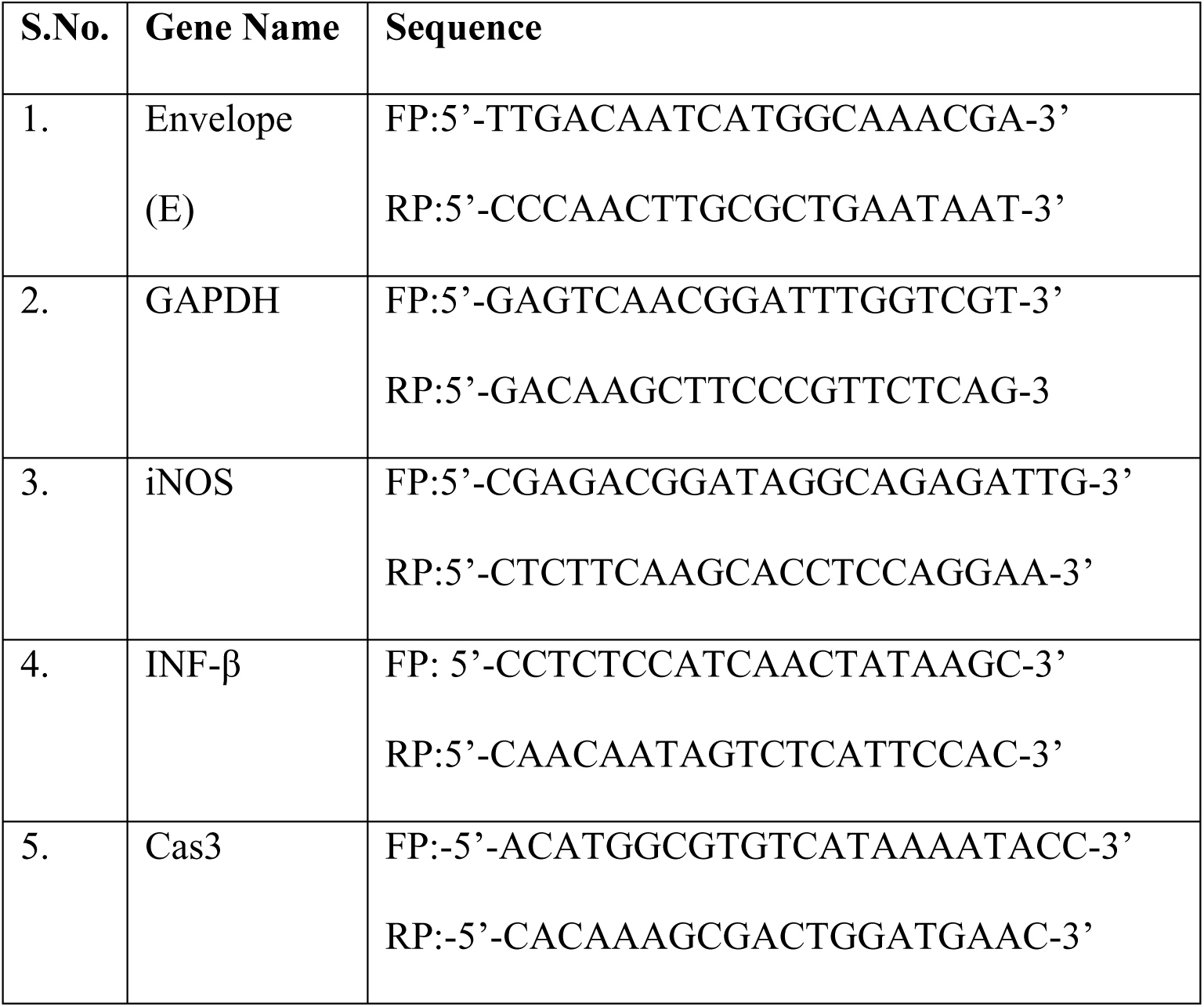
List of primer sequences.

**Supplementary Table No.2:**
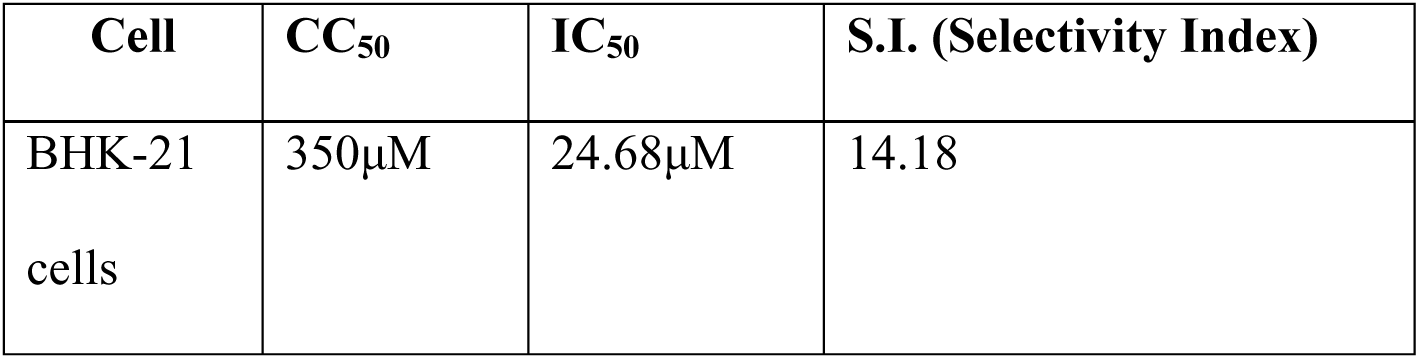
Selectivity Index of TM.

